# Neurogenomic diversity enhances collective antipredator performance in *Drosophila*

**DOI:** 10.1101/2024.03.14.584951

**Authors:** Daiki X. Sato, Yuma Takahashi

## Abstract

Collective behavior is a unique social behavior that plays crucial roles in detecting and avoiding predators. Despite a long history of research on the ecological significance, its neural and genetic underpinnings remain elusive. Here we focus on the mesmerizing nature that visual cues from surrounding conspecifics alleviate the fear response to threatening stimuli in *Drosophila melanogaster*. A large-scale behavioral experiment and genome-wide association analysis utilizing 104 strains with known genomes uncovered the genetic foundation of the emergent behavioral properties of flies. We found genes involved in visual neuron development associated with visual response to conspecifics, and the functional assay confirmed the regulatory significance of lamina neurons. Furthermore, behavioral synchronization combined with interindividual heterogeneity in freezing drove nonadditive, synergistic changes in group performance for predatory avoidance. Our novel approach termed genome-wide higher-level association study (GHAS) identified loci whose within-group genetic diversity potentially contributes to such an emergent effect. Population genetic analysis revealed that selective pressure may favor increased responsiveness to conspecifics, indicating that by-productive genomic diversity within the group leads to a collective phenomenon. This work opens up a new avenue to understand the genomics underpinning the group-level phenotypes and offers an evolutionary perspective on the mechanism of collective behavior.

## Introduction

Group formation is one of the most common behaviors in social animals and plays an essential role in foraging, mating, and avoiding predators^1,2^ and often leads to notable behavioral patterns called collective behavior^3^. From a bird’s-eye view of understanding that assumes virtual access to social information that is not directly available, recent studies have more focused on sensory information processing at the individual level as the emergent mechanism of collective behavior^4–8^. The fruit fly (*Drosophila melanogaster*), a model organism in genetics and neuroscience, may serve as a model system for studying group behavior^9–11^. A significant aggregation of flies emerges through social interaction between individuals, mostly mediated by the sense of visual cues from the surrounding environment^12^. Several research emphasized the functional importance of social aggregation of flies, e.g., flies in a group show less fearful response toward threatening stimuli compared to those in a solitary condition^13,14^. Furthermore, such fear-alleviating “group effect” is strong in a large group of individuals, suggesting that flies have sophisticated social cognitive systems, presumably including the capacity to discriminate numbers^14,15^.

In addition to the neurological mechanisms, there has been growing interest in the genetics underpinning group behavior. The social structures of fruit flies determined by network properties differ greatly among genetically distinct strains^9^ and are probably subject to natural selection^16^. Extensive screening of zebrafish found numerous mutant strains with defects in collective behavior^17,18^, confirming the idea that genetic variation controls interindividual interaction rules. Recent genomic techniques employing genome-wide association studies (GWAS) have also found genetic polymorphisms connected to eusociality in a unique wasp species^19^ and potential loci linked to sociability in guppies^20^. These studies suggested that animals’ intricate group behavior is likely encoded by their genomes and be impacted by specific genetic variants. However, identifying the genomic variants underlying group behavior of non-eusocial insects is difficult due to the paucity of model systems that can associate phenotypes and genotypes at the individual and group levels.

Therefore, the *Drosophila* genetic reference panel (DGPR)^21^, consisting of roughly 200 inbred strains of *D. melanogaster*, is a powerful tool to dissect genomic variations underpinning both individual and group behaviors. These strains are derived from a single population in North Carolina and represent both genotypic and phenotypic variation in the natural environment. The DGRP has been employed in GWAS targeting various physiological and behavioral phenotypes, such as activity levels^22^, aggression^23^, grooming behavior^24^, and circadian rhythms^25^. While these studies demonstrated the utility of the DGRP in the study of behavioral genetics, none focused on the emergent property of group-level traits. The unique feature of the DGRP that the individuals within a strain are considered genetically identical facilitates quantification of group- and individual-level phenotypes linked to the genomic variation. In addition, recent studies have advanced our comprehension of the influence of individual heterogeneity on group dynamics^26–30^. Thus, the complex mechanisms and genetic hallmarks can be understood by assembling mixed groups of genetically diverse strains. Leveraging the genome sequences further facilitates the exploration of the genetic basis of the impacts of such diversity. These techniques shed light on the genomics of collective behavior and offer a fresh way to investigate the genetic underpinnings of higher-level phenotypes transcending individual organisms (Fig. 1a).

**Figure 1.**
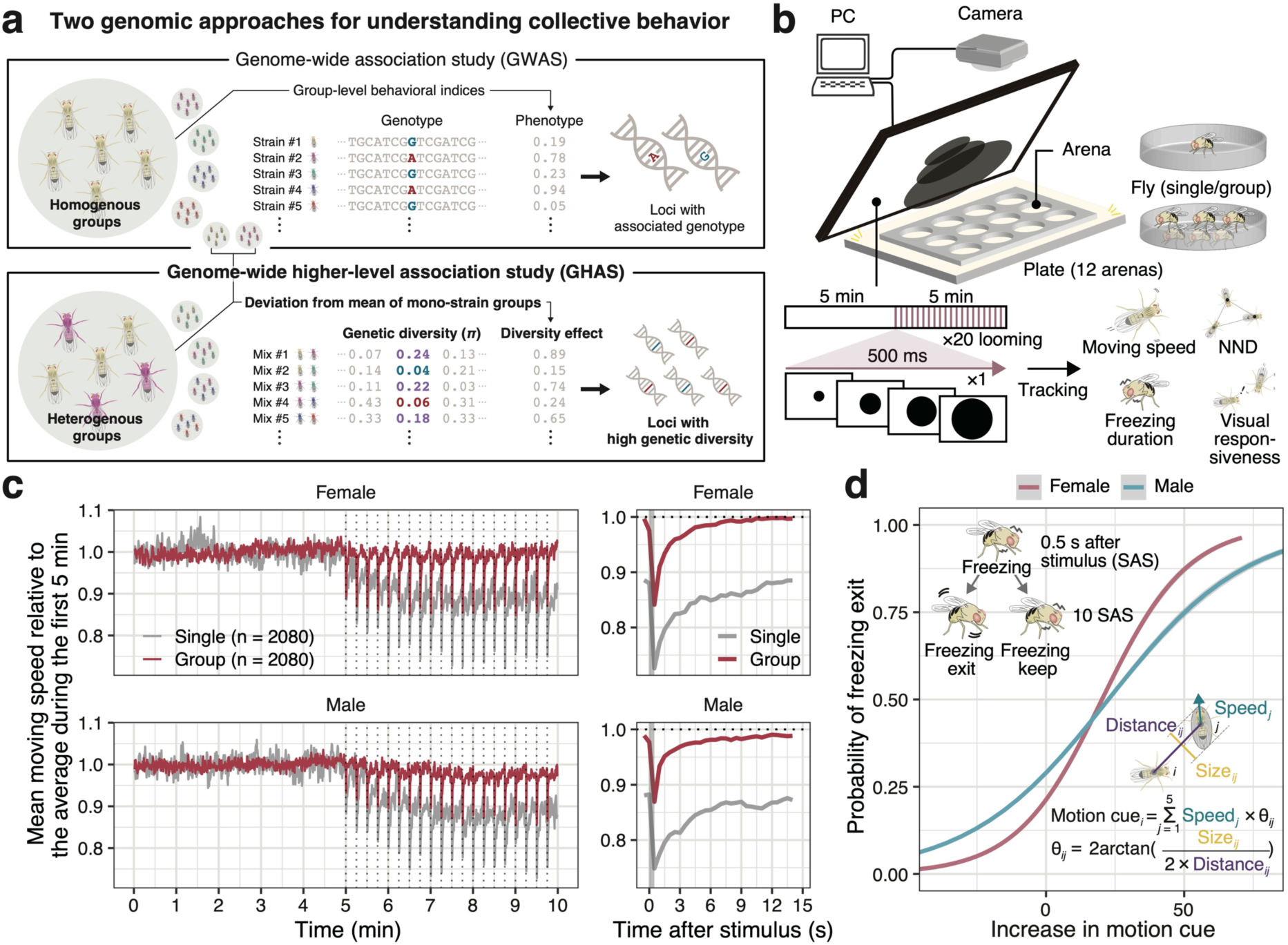
Study scheme, experimental setups of behavioral recordings, and flies’ collective response to threat. **(a)** Two genomic approaches to understand the molecular mechanism of collective behavior are illustrated. Conventional GWAS approach seeks genetic variants whose genotype is associated with a given group-level behavioral trait. Our novel approach, genome-wide higher-level association study (GHAS), detects loci whose genetic diversity in experimental groups is associated with the nonadditive effect of the combination of strains in the groups (called “diversity effect”). **(b)** The experimental setup was adopted from Ferreira and Moita, 2020. Single or grouped flies in an arena were exposed to looming stimuli, and their tracked coordinates were converted to behavioral indices such as moving speed, nearest neighbor distance (NND), freezing probability, and visual responsiveness to other individuals. **(c**) The average moving speed relative to that of the first 5 min across 104 strains. The dotted lines and a grey bar in left and right panel, respectively, indicate the looming stimuli. **(d)** The logistic curve to model the probability of freezing exit as a function of increased conspecific motion cues. The freezing exit was defined for each fly as the proportion of stimulating trials following which it walked > 4 mm/s at 10 s after stopping at 0.5 s. The motion cue was calculated as described in the panel.

The current study sought to clarify genomic loci underlying group behavior of fruit flies, especially under a threatening environment. Combining a large-scale behavioral experiment with GWAS detected genomic loci relative to social responsiveness, which are involved in the development of visual neurons. Additionally, single-cell transcriptomic analysis and mutant behavioral screening demonstrated that differential regulation of lamina neurons influences the visual response to other conspecifics. We also found that genetic diversity among individuals led to nonadditive flexible changes in group behavior, and this emergent impact could be attributed to the loci involved in neuronal development. Taken together, these results show a conceptual breakthrough in the genomics study of group-level behavioral traits and shed light on the genetic and evolutionary mechanisms behind the collective flow of visual information in fruit flies.

## Results

### Genetic variation affects both solitary and group behavioral traits of flies

Flies’ behavior was recorded and monitored for 10 min with repeated looming stimuli (500-ms stimulus for 20 times) displayed within the latter 5 min, as described in previous studies^13,14^ (Fig. 1b). Multiple measures, such as moving speed and head angle averaged at the 0.5-s bin, were used to compute behavioral indices, including locomotor activity, sociality, freezing against fear stimuli, and visual responsiveness to other conspecifics (Fig. 1b; see Methods). The index of baseline locomotor activity was computed using the average moving speed during the first 5 min with no stimuli. A notable difference was seen between strains [*P* < 0.001, linear mixed model (LMM); Fig. S1a], and significant correlations were found between sexes and social circumstances (*r*^2^ = 0.56–0.68 and *P* < 0.001; Fig. S1b), which was consistent with the remarkable hereditary influence on the trait (*H*^2^ = 0.94–0.96). Interestingly, the group effect on the moving speed varied across the sexes, wherein female flies showed reduced while male flies displayed increased moving speed when grouped (Fig. S1b).

Next, we focused on group experiments and evaluated the average nearest neighbor distance (NND) between six flies at rest as a metric of sociality. In our investigation of 104 strains, the majority showed smaller NND compared to the *norpA* mutant strain, which exhibits visual abnormalities, or the pseudo-group data that are generated from random combinations of tracked data taken from individual flies in the solitary condition. Specifically, among females, 98 strains exhibited smaller NND than the *norpA* mutants, and 103 strains showed smaller NND compared to the pseudo-group data (Fig. S2). In males, 72 strains had smaller NND than the *norpA* mutants, and 86 strains exhibited smaller NND compared to the pseudo-group data (Fig. S2). This result confirms that flies are socially aggregated species^12^, and visual information plays an important role in it. In the meantime, several strains were likely to disperse around the arena, suggesting that the genetic variation contained in the DGRP represents a wide spectrum of social phenotypes in a natural population. Despite a prior study suggesting that females are likely to aggregate more than males^12^, the effect of sex was not significant (*P* = 0.91, LMM), while the number of strains showing high levels of sociality (i.e., smaller NND than controls) was larger in females than in males as mentioned above (Fig. S2). Significant effects of strains were detected for NND as well (*P* < 0.001, LMM; *H*^2^ = 0.40 for females and *H*^2^ = 0.53 for males), aligned with the findings from an earlier investigation on genetic impact on the flies’ social network^9,16^.

### Significant decline in locomotor activity by threatening stimuli

We examined the effect of looming stimuli, mimicking threatening objects such as an approaching predator, on flies’ behavior and found a drastic decrease in locomotor activity in response to a stimulus when flies were single, whereas in a group, they quickly returned to the baseline (Fig. 1c). Defining the decrease in moving speed in response to the threatening stimulus as a signature of freezing, we measured and compared the mean length of freezing during the 15 s after the stimulus among strains (see Methods). The outcomes validated the significant effects of strains, social circumstances, and their interactions, except for the items with sex (*P* < 0.001, LMM; *H*^2^ = 0.68–0.84; Fig. S3).

We then investigated the spread of visual information among grouped flies under threat^14^, by modelling the probability of freezing exit as a function of increased conspecific visual motion cues (Fig. 1d) perceived by individual flies within 10 s since a stimulus. The probability of freezing exit logistically increased as the amount of perceived motion cues increased (Fig. 1d), suggesting remarkable conformity in flies’ behavior triggered by visual recognition of movement of nearby flies. Based on the variation in the reaction curve of the conditions (i.e., strains and sexes; Fig. S4a), we defined the intercept of the logistic model as a phenotype representing the threshold of visual reaction toward conspecifics (termed as visual responsiveness hereafter). The significant effects of sex, strains, and their interactions on visual responsiveness (*P* < 0.001 for all items, LMM; Figs. S4b-c) revealed genetic influence on the visual reaction (*H*^2^ = 0.53 for females and *H*^2^ = 0.51 for males). Strikingly, most of the DGRP strains displayed higher levels of visual sensitivity toward conspecifics than the mutant strain (*norpA*) (Fig. S4b), validating the difference in their visual responsiveness.

### Genomic loci underlying social phenotypes of flies

Next, we conducted genome-wide association analyses (GWAA) independently for both sexes to evaluate each of the three phenotypes: freezing duration, NND, and visual responsiveness. The GWAA for freezing duration was performed independently for solitary and grouped conditions. The genetic distances between strains showed that the used 104 strains were genetically diverse enough to cover the most of 205 DGRP strains (Fig. S5a). Prior to the GWAA, we estimated the single nucleotide polymorphism (SNP)-based heritability and the genetic correlations between the above phenotypes. The SNP-based heritability estimates were overall lower than the corresponding *H*^2^ estimated from phenotypic variance among strains (Fig. S5b). Although similar, the genetic correlations were greater than the phenotypic relationships (Figs. S5c-d), possibly indicating that a small number of loci were accountable for the phenotypic variations. The GWAA results are displayed in Manhattan plots, whereas the number of genes adjacently located to the associated loci (*P* < 1.0 × 10^−5^) is shown in Venn diagrams (Fig. S6). Here we focus on the results of visual responsiveness, where multiple significantly associated loci stood out (Fig. 2a). Gene set enrichment analysis (GSEA) was used to examine the functional enrichment of detected genes, and significant gene ontology (GO) terms, including “eye development,” “axon guidance,” or “neuron projection,” were identified (Fig. 2b). Additionally, two genes (*kirre* and *Ptp99A*) were found to be associated with the trait in both sexes (Fig. 2c), and a commonly detected SNP in *Ptp99A* (3R:25292746) revealed a clear genotypic effect (Fig. 2d).

**Figure 2.**
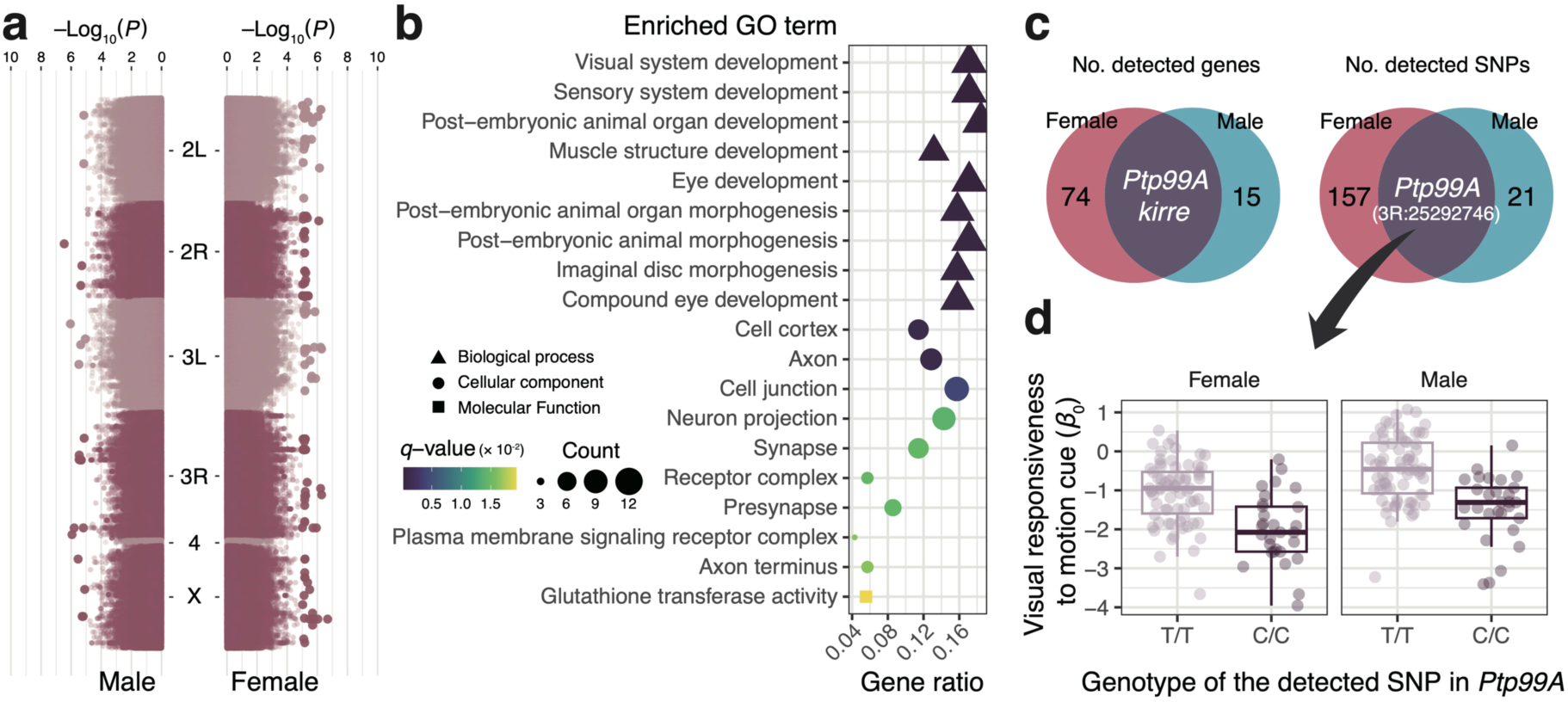
Genome-wide association analysis (GWAA) suggests *Ptp99A* as a candidate gene underlying collective behavior of flies. **(a)** Manhattan plots showing associated loci with visual responsiveness across genome. Colors represent different chromosomes, and the size of points indicates statistical significance (*P* < 1.0 × 10^−5^). **(b)** Gene ontology (GO) enrichment analysis revealed that many of genes detected in GWAA were involved in the development of visual neurons. **(c)** Venn diagram showing the overlap between detected genes in males and females. **(d)** An intronic variant (3R: 25292746) in *Ptp99A* detected in both sexes was a top candidate possibly regulating visual responsiveness to other conspecifics.

### Single-cell transcriptomics reveals that *Ptp99A* SNP affects its expression in lamina neurons

The two candidate genes, *kirre* and *Ptp99A*, are known to be cell surface molecules (CSMs) and modulate the extension of lamina (L1–L5) neurons^31,32^; therefore, we investigated a publicly available single-cell RNA-sequencing (RNA-seq) dataset of the developing visual neurons of *D. melanogaster*^32^. The dataset contained gene expression data of *W*^1118^ flies crossed with DGRP strains, rendering it useful in examining the impact of the identified variant on the expression pattern of *Ptp99A*. Based on the annotation of neuronal cells (24–96 h after pupal formation)^32^, we found *Ptp99A* expression in nearly all types of visual neurons; nonetheless, clear expression was detected in the L5 neuron (Fig. 3a). Additionally, differential expression was observed between the genotypes (C and T alleles) at various developmental time points in L1, L4, and L5 neurons particularly at later stages (Fig. 3b), indicating the regulatory significance of this variant.

**Figure 3.**
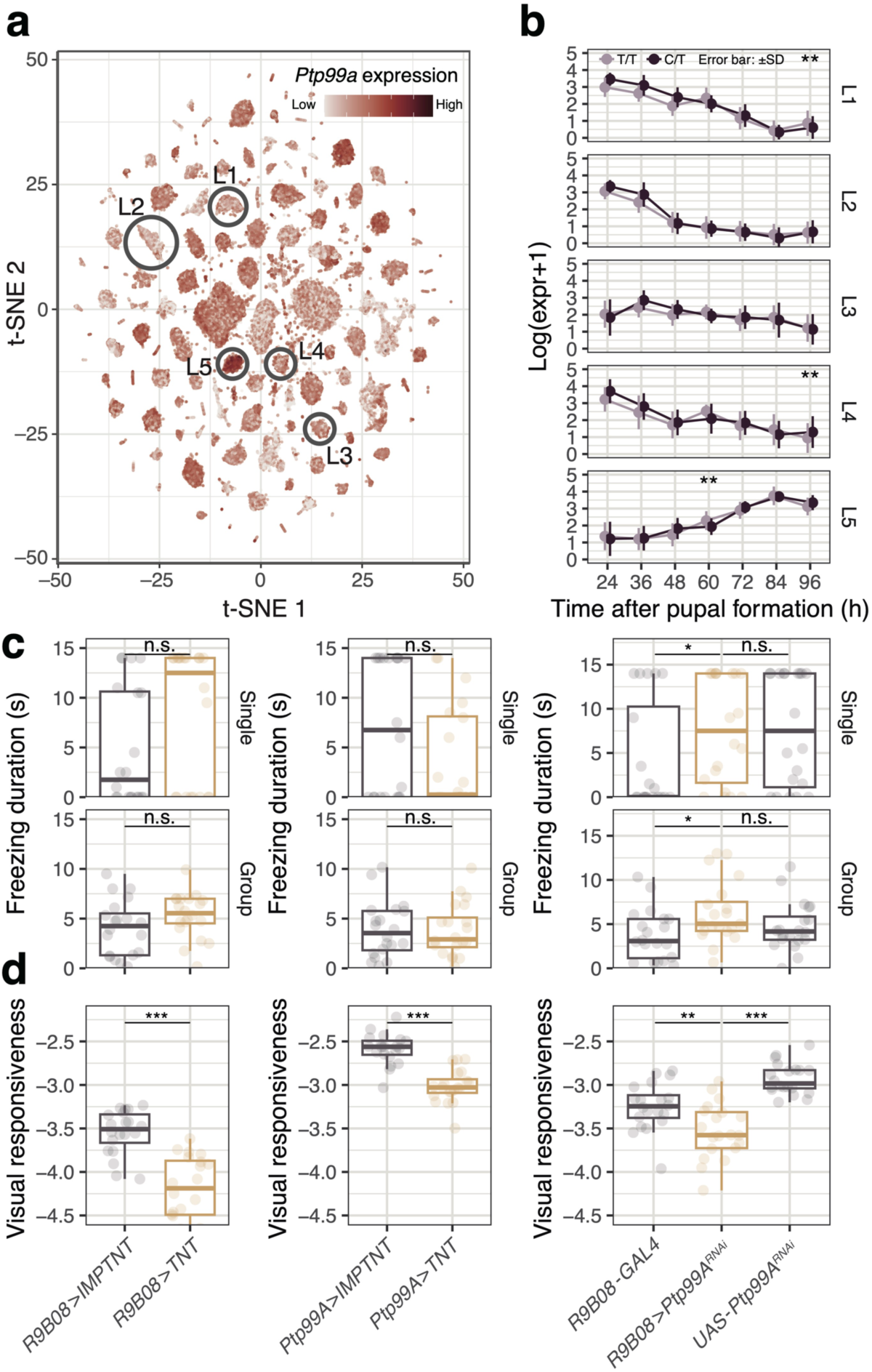
Transcriptomic and neural evidence indicate the importance of lamina neurons in interindividual interaction. **(a)** Expression pattern of *Ptp99A* in *Drosophila* visual neurons visualized on t-SNE dimensions. Lamina (L1–L5) neurons are indicated in circle. Cell annotations followed the results of clustering described in Kurmangaliev et al. 2020. **(b)** Expression pattern of *Ptp99A* in L1–L5 neurons and its difference between the genotype (3R:25292746). Statistical significance was evaluated by GLMM assuming Gamma distribution with strain and replicate as random effects. **(c)** Freezing duration of mutant strains for both solitary and group conditions are shown in boxplots. **(d)** Visual responsiveness of mutant strains is shown in boxplots. *P*-value was calculated by Wilcoxon’s rank sum test for **(c)** and **(d)** (n = 20 for each group). \*\*\**P* < 0.001, ** 0.001 ≦ *P* < 0.01, *0.01 ≦ *P* < 0.05.

### Silencing lamina neurons causes defects in the visual response to conspecifics

To further validate the functionality of lamina neurons and the regulation of candidate genes in motion detection of conspecifics, we used the GAL4/UAS system and inhibited the activities of certain neurons with tetanus toxin light chain (TNT). Silencing lamina neurons using the pan-lamina neuron driver *R9B08*-*GAL4* with *UAS*-*TNT* markedly decreased visual responsiveness, although freezing level stayed constant (Fig. 3c). Next, we investigated the consequences of *Ptp99A* knockdown (KD) in lamina neurons using *UAS*-*Ptp99A^RNAi^*. Similar to the results of *R9B08*-*GAL4*>*UAS*-*TNT*, the conditional KD (*R9B08*-*GAL4*>*UAS*-*Ptp99A^RNAi^*) resulted in the reduced level of visual responsiveness. Silencing the *Ptp99A*-expressing neurons (*Ptp99A*-*GAL4*>*UAS*-*TNT*) also significantly decreased the responsiveness, suggesting that *Ptp99A* may affect the development of lamina neurons, leading to the changes in optical characteristics and social behavior.

### Genetic heterogeneity within a group enhances flexible changes in freezing behavior

We examined the effect of genetic diversity within a fly group on the antipredator performance. With 15 genomically diverse strains (Fig. S5a), we created groups of 6 female flies by combining 3 individuals from each of two different strains and conducted behavioral experiments across a total of 105 strain combinations. The measured indices of the mixed-strain group were compared to the expected values, which were calculated as the mean of measured values from the groups of each strain that constitute the mixed group. Interestingly, the moving speed was dramatically decreased just after the visual stimulus in mixed-strain groups (P < 0.001 for 0.5–1.5 s after stimulus, Wilcoxon’s signed rank test with Bonferroni correction; Figs. 4a and S7). Reduced locomotor activity is the optimal evasion tactic under high predatory pressure^33,34^; the observed changes in behavior may thus be adaptive. Therefore, we computed the virtual fitness values that incorporate the balance between vigilance and exploration based on the time-series changes in moving speed (Fig. 4b; see Methods); the fitness values were significantly greater in mixed-strain groups (P < 0.001, Wilcoxon’s signed rank test; Fig. 4c). The difference between the expected and observed values of virtual fitness in mixed-strain group was termed as the “diversity effect” (Fig. 4b). Most combinations of strains displayed a positive diversity effect (i.e., overyielding), with several showing higher fitness values than the maximum of the two strains comprising the given group (i.e., transgressive overyielding) (Fig. S8). Furthermore, we discovered that the difference in freezing duration between the two strains was positively correlated with the diversity effect (Fig. 4d), indicating that behavioral synchronization among individuals with varying freezing parameters expedited the response to visual threat.

**Figure 4.**
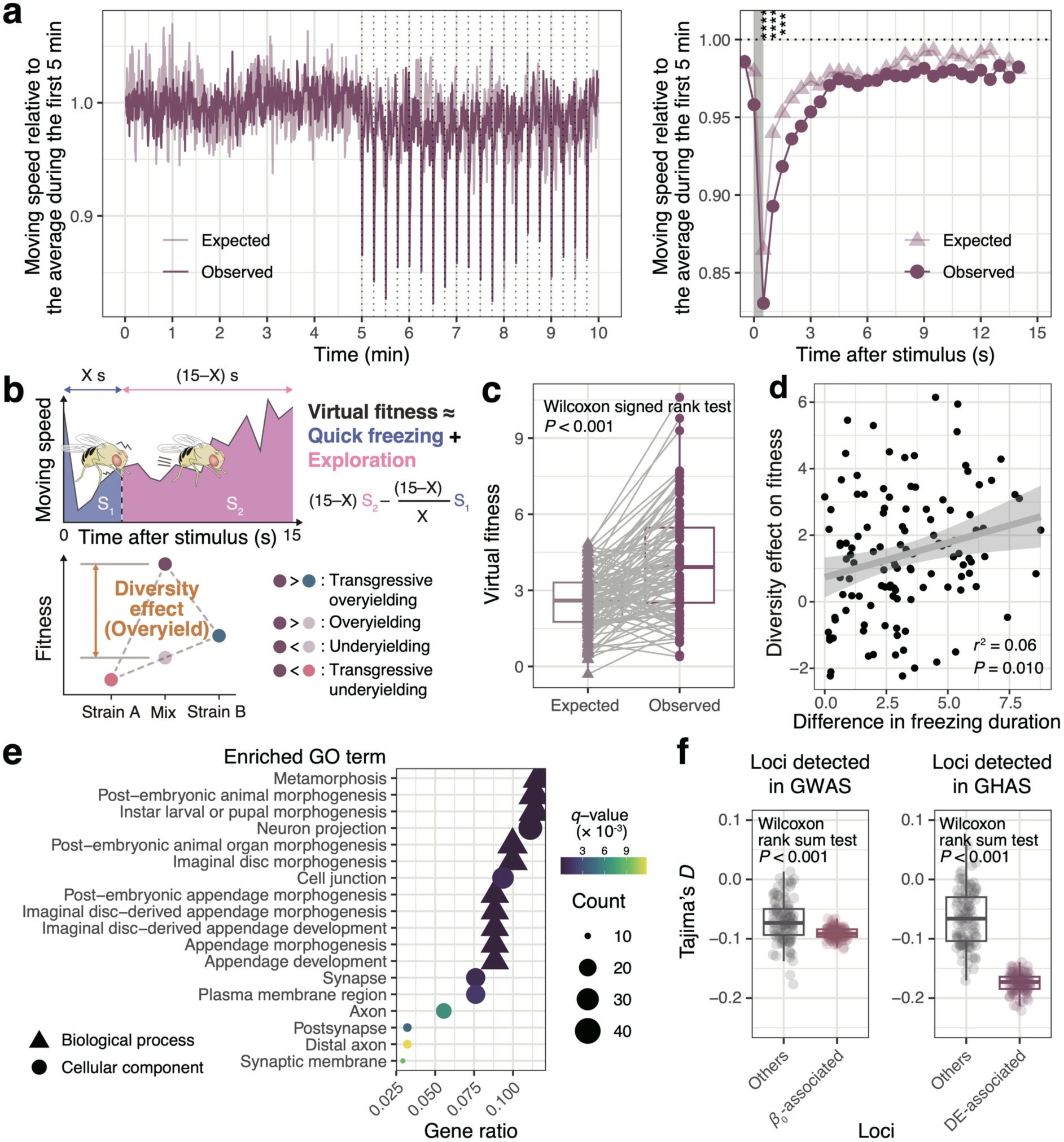
Genetic heterogeneity within group results in nonadditive changes in group behavioral dynamics. **(a)** The mean relative moving speed was significantly lower in mixed-strain groups immediately after the stimulus, suggesting that behavioral heterogeneity enhanced freezing. The exposed timing of stimulus is indicated in a grey bar. Asterisks denote statistical significance, calculated by Wilcoxon’s signed rank test with Bonferroni correction (n = 105 for each group); \*\*\*\**P* < 0.0001, *** 0.0001 ≦ *P* < 0.001. **(b)** Schematic illustration of calculation of virtual fitness. Assuming that freezing under predatory pressure and exploration otherwise would be adaptive, we calculated the weighted sum of moving speed, in which the coefficient was negative in the freezing term (colored in blue) and positive in the exploration term (colored in pink). We defined the diversity effect as the deviation from the mean virtual fitness of mono-strain groups, representing the nonadditive increase in fitness by genetic and/or behavioral diversity in fly groups. **(c)** The mixed-strain groups showed significantly high levels of virtual fitness. A grey line connected to two points represents a strain, and statistical significance was evaluated by Wilcoxon’s signed rank test (n = 105 for each group). **(d)** Correlation of the difference in freezing duration between two strains and the diversity effect in virtual fitness of their mixture (n = 105). **(e)** The results of GO enrichment analysis of loci detected in genome-wide higher-level association analysis (GHAS). **(f)** Tajima’s *D* was significantly lower in loci detected in both GWAS for visual responsiveness and GHAS. The *P*-value was calculated by Wilcoxon’s rank sum test (n = 100 for each group).

Given that the diversity effect on virtual fitness is a group-level phenotype, we used GWAA to pinpoint the genomic loci linked to with the phenomenon. Our approach, termed genome-wide higher-level association study (GHAS), addresses higher-level (i.e., group, community, or population) phenotype as well as genetic profile determined by population genetic statistics calculated for a particular set of individuals (Fig. 1a). In the present study, we calculated the nucleotide diversity (*π*) as a metric of genetic diversity for 105 pairs of strains across the genome and found loci whose nucleotide diversities were positively correlated with the diversity effect. The top associated loci were those related to metamorphosis, organ morphogenesis, neuron projection, and axons (Fig. 4e). Remarkably, the genes adjacently located to the identified loci included many of those found in GWAS for the visual responsiveness, including *kirre* and *Ptp99A* (Fig. S9).

The genetic diversity that confers benefits at the group level may be maintained by natural selection such as balancing selection. Therefore, we investigated how the selection acted on the loci detected in GWAS for visual responsiveness and GHAS. The findings showed that compared to other loci in the genome, the identified loci in both analyses had a negative skew in Tajima’s *D* values (Fig. 4f), suggesting directional selection on these loci throughout evolution.

### Agent-based simulation supports simple mechanisms of collective freezing

Finally, we performed computer simulations using an individual-based model (IBM) to test the behavioral mechanisms behind collective freezing and its diversity effect found in this work. We replicated the identical experimental setup and placed either one or six agents in the simulation. The moving speed of the agent at time *t* (*V*_*t*_) followed a Lévy distribution with a mean speed of 3.8 mm/s, derived from our experimental data of female flies, and a shape parameter *α* = 2, with a change in moving direction (*D*_*t*_) following a normal distribution with a mean of 0 and a standard deviation of *π* (Fig. 5a). Furthermore, we replicated the agents’ reactions to repeated visual stimuli by characterizing the decrease in moving speed. Under group conditions, the changes in surrounding motion cues over time were incorporated into their own velocities to simulate synchronization in moving speed with other individuals; the result showed a notable difference in the moving speed under stimuli between social circumstances (Fig. 5b), which was consistent with the observation in real flies (Fig. 5b). Moreover, by modifying the parameters related to freezing to represent genetically distinct strains (Fig. 5a), we successfully replicated nonadditive changes in behavior seen in groups consisting of different strains (Fig. 5c). These findings confirmed that the behavioral synchronization via visual information produces collective freezing and that behavioral heterogeneity among individuals creates additional benefits in collective performance.

**Figure 5.**
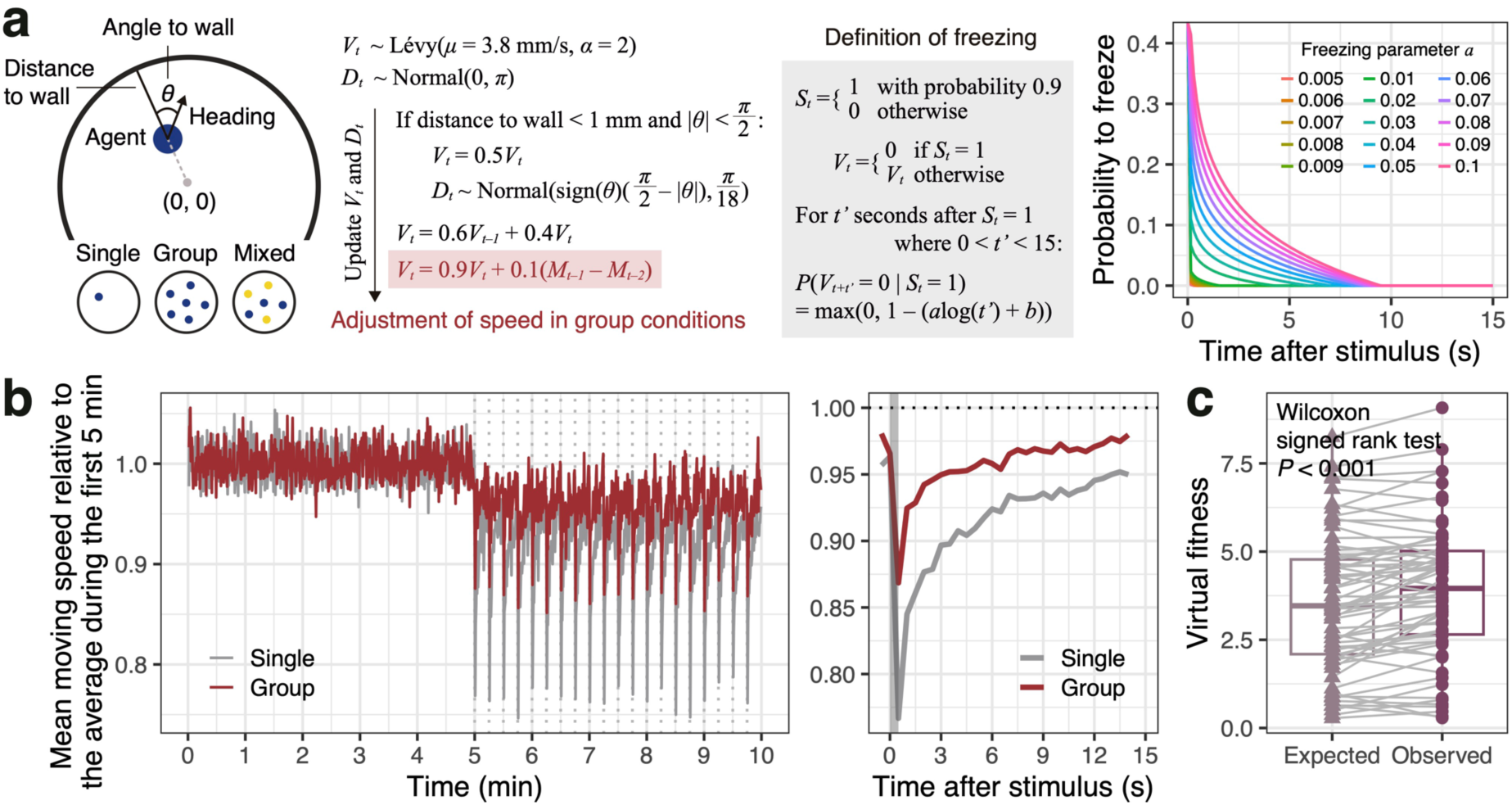
Agent-based simulation replicates the collective behavior enabled by simple mechanisms. **(a)** The parameter settings of agent-based simulation. Single and group models only differ in the speed adjustment mechanism (colored in red) where the change in motion cues of other individuals between previous time steps (*M*_*t*−1_ − *M*_*t*−2_) was incorporated into current velocity (*V*_*t*_). We also reproduced the 15 strains with phenotypic diversity in freezing parameters and simulated mixed-strain group experiments by introducing 3 individuals each from two strains. **(b)** The average moving speed showed a similar pattern to experimental dataset (Fig.1c), strongly implicating that visual response to other individuals change the dynamics of collective freezing. **(c)** The increase in diversity effect on virtual fitness was also observed in simulated dataset, supporting that phenotypic diversity in freezing response could result in flexible changes in group behavior. The statistical significance was evaluated by Wilcoxon’s signed rank test (n = 105 for each group).

## Discussion

While the mechanism underlying collective behavior has long been debated in the field of behavioral ecology and/or physics, the knowledge about its genetic substrates has been limited especially in non-eusocial insects^9,12,16^. In this study, we concentrated on the transmission of visual information among individuals under predator-mimicking stimuli and identified genes linked to collective behavior in flies. The flies displayed freezing in response to looming stimuli, but the degree of freezing reduced upon detecting motion cues from conspecifics under group conditions (Fig. 1). We detected the genetic variants associated with this group effect using GWAS (Fig. 2) and found the candidate genes implicated in the development of visual neurons, such as *kirre* and *Ptp99A*; these are CSMs controlling the interactions among lamina neurons^31,32^. The single-cell transcriptomic data of *Drosophila* visual neurons^32^ showed genotype-dependent differential expression of *Ptp99A* (Fig. 3b). Functional investigation with the GAL4/UAS system additionally revealed that the KD of *Ptp99A* and silencing lamina neurons influence responsiveness to other individuals (Fig. 3c). These results indicated a critical role of visual, particularly lamina, neurons in the collective behavior of fruit flies.

The lamina is a component of the visual system of the fruit fly, serving as the initial layer of visual processing in the compound eye. This processing facilitates the detection of changes in light intensity, moving objects, and color discrimination. The detection of moving objects is one of the key functions occurring in the lamina^35,36^. Lamina neurons include L1– L5 monopolar cells, each of which projects to different layers in the outer medulla. L1–L3 neurons receive photoreceptor signals from R1–R6 neurons in the retina, while L4 and L5 neurons receive indirect input of photoreceptor signals through L2 and L1 neurons, respectively^37,38^. The photoreceptor signals are divided into two parallel motion circuits, ON and OFF pathways^35,36^, that send luminance signal increase and decrease, respectively, playing an essential role in edge detection and contrast enhancement^38,39^. Herein, we observed differential expression of *Ptp99A* in several lamina neurons with the 3R:25292746 genotype at various developmental stages (Fig. 3b). Although the neural mechanism is yet unclear, such genotypic difference might cause developmental disparities in the lamina neurons. This may result in differential processing of optical information as well as visual responsiveness to moving conspecifics and, consequently, in emergent dynamics of the group behavior, potentially in fitness.

The heterogeneity of behaviors among individuals and its effect on group-level characteristics are currently under intensive focus^26–30^. In this work, we introduced an equal number of individuals with various genetic backgrounds into a group to test the effect of behavioral heterogeneity on collective freezing through interindividual interactions. The findings showed that in mixed-strain groups, the degree of freezing, immediately following a stimulus, was higher than expected (Fig. 4a), suggesting that heterogeneity in behavior within a group produces nonadditive changes in the behavioral performance at the group level. Additionally, we established an analytical framework, GHAS (Fig. 1a), and identified loci where genetic diversity within the group is associated with an increase in virtual fitness. Remarkably, the detected loci were overlapped with those found to be associated with visual responsiveness in GWAS and included many genes involved in the development of the visual neurons, including *Ptp99A* (Fig. S9). This result suggested that phenotypic diversity in the visual function among individuals underlies the adaptive changes in group behavior. Notably, the differences in the duration of freezing but not visual responsiveness to other conspecifics between strains were positively correlated with the diversity effect in mixed-strain groups (Figs. 4d and S10), indicating that behavioral heterogeneity and adequate synchronization among individuals led to emergent behavioral changes at the group level. Furthermore, simulations utilizing an IBM showed that conformity to the motion cues of other individuals decreased the group-level freezing and that the interindividual difference in freezing parameters induced adaptive behavioral changes in the group (Fig. 5c). These results clarify that the emergent phenomena observed in this study occur through straightforward interaction mechanisms.

Furthermore, we explored the selective pressures involving group behavior. In this work, we examined the freezing response to visual stimuli and found that visual responses to other conspecifics might lead to emergent changes in group behavior. The behavioral heterogeneity and conformity among individuals in groups composed of strains with varied behavioral features resulted in nonadditive behavioral changes at the group level, potentially increasing fitness. However, population genetic analysis using Tajima’s *D* showed that the genetic diversity associated with this diversity effect might be under directional selection rather than balancing selection (Fig. 4f), suggesting that the genetic diversity at these loci may have a detrimental effect on individual fitness. Based on the observation of *Ptp99A* found by GWAS, strains carrying the major T allele showed higher responsiveness to motion cues (Fig. 2d), whereas mutants with inhibited neural transmission in the lamina showed lower visual responsiveness (Fig. 3c). These results indicated that selective pressure might enhance the responsiveness to conspecifics. Consequently, the diversity effect brought about by genetic diversity among individuals on group behavior might be a passing phenomenon, possibly differing from fitness on the evolutionary timeline. Thus, our results do not merely disprove group selection but also offer a new direction of research by demonstrating its verifiability from a genomic perspective.

Finally, our findings provide an insight into the functional roles of neuronal diversity and its evolutionary importance. Synaptic and non-synaptic partners of lamina neurons express several couples of interacting CSMs^32^, including *Ptp99A* and *kirre*. This redundancy helps functional compensation of CSMs, reducing the negative effects of their disruption^40,41^. Moreover, this may also amplify the developmental robustness of hardwired lamina neurons that were shown to specifically converge to the lamina precursor cell cluster, unlike other types of neurons^42^. Taken together, our data demonstrated directional selection on loci involved in CSMs, leading to increased visual reactivity to conspecific motions. Such selective pressures may decrease the neuronal diversity of lamina neurons, contributing instead to their robustness. However, the effect of these factors on the general development of the brain and its neuronal diversity remains unclear, providing an important direction for future study.

## Methods

### Fly strains and keeping conditions

A cohort of 104 strains derived from the *Drosophila* Genetic Reference Panel (DGRP) was used in our experiment. This panel consists of 205 inbred and genetically distinct strains of *D. melanogaster*, representing natural genetic diversity within a population captured in North Carolina, US. Flies were kept at 25 °C with about 40–60% humidity in a 12L:12D cycle, with standard food (Bloomington Formulation, Nutri-Fly™, #66-113, Genesee Scientific). A visually defective mutant strain, *w*[*] *norpA*[P12]^43^, was used to test the behavior of blind flies and the contribution of visual information in our assay. The following lines were used for functional assay of lamina neurons and *Ptp99A*: *R9B08*-*GAL4*, *Ptp99A*-*GAL4*, *UAS*-*TNT*, *UAS*-*IMPTNT*, *UAS*-*Ptp99A^RNAi^*. All fly strains were obtained from the Bloomington *Drosophila* Stock Center (BDSC), and the list of used strains with the stock number is summarized in Table S1. Adult flies of both sexes aged at 2–4 days since eclosion were tested in the first behavioral screening with DGRP strains and *norpA* mutant (n = 20 per sex and social circumstance for each strain). Given the observed chasing behavior among male flies in some strains, which likely impacts their social interaction, we used only female flies in a subsequent mixed-strain group experiment (n = 10 for each pair of strains) and mutant assay (n = 20 for each condition of strains).

### Behavioral assay under looming stimuli

Behavioral assays were conducted during a Zeitgeber time of 0–12 within an incubator keeping the same conditions of temperature and humidity as housing. To avoid sexual behavior which can disturb response to threatening stimuli and conspecifics, we used single-sex groups of flies for experiments. The experimental apparatus was set as described in Fig. 1b referring to previous studies^13,14^. Briefly, either single or grouped (n = 6) flies were anesthetized with CO2 and placed in an arena with sloped walls (30 mm in diameter and 2 mm deep), where flies were restricted from flying. After 30 min of acclimation, including the last 5 min being recorded, they were exposed to looming stimuli for 5 min (20 trials of a black circle expanding within 500 ms on a white background every 15 s) as previously described^13^. The stimulus was presented at 60 Hz on a 13.3-inch EVICIV monitor, tilted 45° toward the experimental plate. The plate had 12 arenas and an LED board (147 mm width × 115 mm height, 7500 lx, TLB-MP, Asone Co., Japan) placed under it. Each condition (strain, sex, and social circumstance) of flies was randomly placed in one of the arenas, and the behavior was recorded at 1920 width × 1080 height in pixel resolution and approximately 50 frames per second using a USB3 camera (DMK33UX290, The Imaging Source, Germany) controlled by a custom python script. Recorded videos were trimmed for each arena, and flies’ behavior was primarily tracked by the tracktor software^44^, followed by our custom script to fix tracking errors. To further reduce the calibration errors of tracking, the weighted moving average of coordinates over a time span (10 preceding time points including a time point of interest) was used for analysis. The coordinates were subsequently used to calculate the distance traveled from a previous time point, speed (traveled distance divided by frame intervals), and head angle (calculated by the movement between time points 50 ahead and behind the time point of interest) for each time point. These measures were averaged at 0.5-s intervals and used for subsequent analyses. Locomotor behavior was briefly dichotomized as walking when their speed was > 4 mm/s or stopping otherwise^13^. Freezing was defined when a fly that stopped 0.5 s after a stimulus continued to stop 10 s after the stimulus, otherwise referred to as freezing exit.

### Calculation and analysis of behavioral indices

The average moving speed during the first 5 min with no stimuli was assessed to be an index of baseline locomotor activity. To quantify the level of the sociality of flies, we calculated the NND at a time point when all the flies in a given arena were expected to be at rest (i.e., a time point when the average moving speed across flies were minimum during the 5 min). For comparison, pseudo-group data consisting of random combinations of single fly data were created and evaluated as a control with no social preference. In this process, an experimental date, time, and used plate and arena was randomly chosen from the information of constituting flies and used as covariates for the data. In the latter 5 min of the experiment, the average time taken to restore the moving speed to the baseline level (i.e., the mean moving speed during the first 5 min) after exposure to stimuli was used as an indicator of the fear response to threatening stimuli. In order to quantify the flies’ response toward other individuals, we first calculated the “motion cue” that each fly is expected to perceive from other flies, defined as follows (Fig. 1d; modified from Ferreita and Moita, 2020):

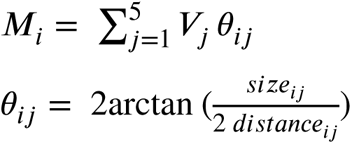

where for a focal fly *i* in a group, the motion cue it perceived (*M*_*i*_) was defined as the sum of other flies’ speed multiplied by the visual angle. *V*_*j*_and *θ*_*ij*_ represent the moving speed and the visual angle of the individual *j* on the retina of the individual *i*, respectively. The distance between individuals *i* and *j* (*distance*_*ij*_) was calculated from their centroids, and the visual size of the individual *j* to *i* (*size*_*ij*_) was calculated by projecting the individual *j*’s body (represented by an ellipse with 1 and 2 mm in a smaller and larger radius, respectively) on the body axis of the individual *i* as described in Fig. 1d. Next, we modeled the probability of freezing exit (*P*_FE_) explained by the increase in perceived motion cues in 10 s after a stimulus with a following logistic regression:

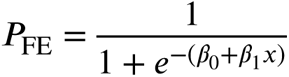

Here, the intercept *β*_0_was used as an index of response threshold, representing flies’ visual responsiveness toward motion cues of other conspecifics. To accurately estimate the logistic curve fitting to the visual response of each strain of flies, we pooled the data of all stimulating trials (6 individuals × 20 trials × 20 replicates = 2400 data points at maximum) and conducted above regression for each strain. In case for the dataset of GAL4/UAS strains, we randomly sampled 500 data points from all the dataset of a given strain to perform logistic regression and repeated it for 20 times to obtain the *β*0 estimates for the population.

### Statistical analysis for behavioral traits

Statistical analysis of behavioral data was performed using R 4.1.2 with the *car*, *lmerTest*, and *lme4* packages when necessary. Effects of strains, sex, social circumstances, and their interactions on behavioral measures were assessed with the LMM with experimental date, time, used plate and arena assumed to be random effects. Generalized linear mixed model (GLMM) assuming binomial distribution was used to evaluate the logistic model for visual responsiveness described in Fig. 1d. The broad-sense heritability of traits was estimated by the formula described in previous studies^45,46^ using mean squares for genotype (strain) and environment (residual error) obtained in analysis from variance (ANOVA).

### Genome-wide association analysis

We first performed principal component analysis (PCA) to capture the genetic distance among strains, using a whole set of genotypes of 4,438,427 SNPs for 205 strains obtained from the DGRP2 website (http://dgrp2.gnets.ncsu.edu) with PLINK v1.9^47^. Quality control of SNPs was carried out with conditions: (1) minor allele frequency > 0.01 and (2) SNP call rate > 0.9, leaving 2,672,211 SNPs for subsequent analyses. Prior to GWAA, SNP-based heritability and genetic correlation among phenotypes were estimated by SCORE^48^. We then performed GWAA to identify loci responsible for the three phenotypes (i.e., freezing duration, NND, and visual responsiveness) of flies. The behavioral phenotypes were regressed with the top 10 PCs as covariates using GCTA v1.94.0 with --*fastGWA-mlm* flag^49^, based on a linear mixed model and a SNP-derived sparse genetic relationship matrix. The association analysis was performed independently for each sex, and loci with the *P*-values below the genome-wide significance level as *P* = 1.0 × 10^−5^ were detected to be candidates associated with the given phenotype. GO enrichment analysis for candidate loci was conducted independently for both sexes using the *clusterProfiler* package in R.

### Single-cell transcriptomic analysis of *Drosophila* visual neurons

Single-cell RNA-seq dataset of the developing visual neurons of *D. melanogaster*^32^ was reanalyzed. The dataset included expression data of visual neurons of *W*^1118^ crossed with various DGRP strains at 9 developmental stages (0–96 h APF). Given the substantial difference in tissue composition and coverage^32^, we used the main dataset comprised of 24– 96 h APF in the current study. Each cell was annotated according to the results of the previous study^32^, and t-distributed stochastic neighbor embedding (t-SNE) was used to visualize the clusters of them. The expression levels of *Ptp99A* between the genotype (3R:25292746) detected in GWAS was compared by generalized linear mixed model assuming gamma distribution with strains and replication as random effects using glmer() function implemented in the *lme4* package in R.

### Mixed-strain group experiments and calculation of diversity effect

Among the 104 strains used in GWAS, we selected genomically diverse 15 strains (cf., Fig. S5a) and prepared groups of 6 female flies, composed of 3 individuals from each of the two different strains (i.e., 105 pairs of strain in total; n = 10 for each). Following the behavioral experiment, the average moving speed among individuals relative to that of first 5min in a mixed-strain group was compared with the average values of the two strains that constitute the group. The difference between the observed value of the mixed-strain group and the expected value (namely, the average values of the single-strain groups of the two strains) was defined as the diversity effect (Fig. 4b). Subsequently, considering visual stimuli as hypothetical predators, virtual fitness was defined based on the flexible behavioral changes observed after the visual stimulus. Assuming that freezing under predatory pressure and exploration otherwise would be adaptive, we calculated the weighted sum of moving speed, in which the coefficient was negative in the freezing term and positive in the exploration term (Fig. 4b). Given that the moving speed of flies almost returned to the baseline within 3 s after a visual stimulus, the freezing term was set to 0–2.5 s after stimulus (SAS) and exploration terms was set to 3–14 SAS. Weights were applied to adjust for the duration of each time period in the calculation (Fig. 4b).

### Identifying responsible loci for diversity effect with GHAS

The diversity effect was computed per combination of strains and may be explained by the genetic differences (diversity) between the two strains. To comprehensively explore the responsible loci for the diversity effect, we calculated the nucleotide diversity (*π*) at each locus (1000 bp windows) from the genome sequences of the two strains constituting the mixed group using vcftools^50^ and conducted an association analysis with the diversity effect, which we termed the genome-wide higher-level association study (GHAS). Genetic distances between strains on the PC plane shown in Fig. S5a were used as covariates. Loci with *P* < 8.7 × 10^−6^, corresponding to the Bonferroni-corrected family-wise error rate (α = 0.05 divided by the number of analyzed loci) were identified as candidate loci, and GO enrichment analysis was performed using the R package *clusterProfiler*.

### Test for selection using Tajima’s *D* statistics

If the diversity effect occurring at the group level has a positive impact on actual fitness, it can be inferred that natural selection, specifically balancing selection, is acting to actively maintain genetic variation. To verify this, we used Tajima’s *D*, a population genetic statistic which indicates balancing selection with a positive deviation and directional selection with a negative deviation, relative to the genome average. Thus, we calculated Tajima’s *D* for the loci detected by GHAS and compared the values with those calculated from randomly sampled loci across genome. We applied a permutation test in this analysis, wherein a subset of the detected loci (i.e., 400 out of the 458 loci detected in GHAS) was extracted, and the average Tajima’s *D* was calculated. Likewise, an equal number of loci were randomly selected from the entire genome, and their average Tajima’s *D* was calculated. We repeated this process for 100 times and compared the values between the two groups. Additionally, the same calculations were conducted for loci (1000 bp windows containing the detected variants) associated with visual responsiveness detected by GWAS, to examine the mode of selection.

### Behavioral simulation with individual-based model

To explore the mechanisms behind the observed changes in fear responses in fly groups and the diversity effect in mixed-strain groups, we conducted behavioral simulations using an IBM (Fig. 5a). The arena size was set to 30 mm, and the number of agents in the arena was set to one in the single condition and six in the group and mixed-strain group conditions. Like the actual experiments, each time step was set to 0.02 s, replicating a 10-min experiment (i.e., 30,000 frames in total). The moving speed of the agent at time *t* (*V*_*t*_) followed a Lévy distribution with a mean speed of 3.8 mm/s, obtained from our experimental data, and a shape parameter *α* = 2, with a change in moving direction (*D*_*t*_) following a normal distribution with a mean of 0 and a standard deviation of *π*:

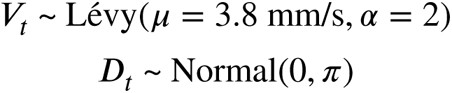

The Lévy distribution was chosen to simulate the natural movement patterns observed in biological organisms. Let the vector representing the agent’s heading direction be 𝕧, and the normal vector to the nearest wall from the agent’s current position be 𝕟. The angle between these two vectors, representing agent’s angle to wall, can be calculated as follows:

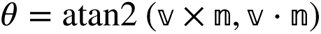

where 𝕧 × 𝕟 represents the z component of cross product of 𝕧 and 𝕟, calculated as *v*_*x*_*n*_*y*_ − *v*_*y*_*n*_*x*_, and 𝕧 ⋅ 𝕟 represents the dot product of the two vectors. If the agent approaches the wall, it decelerates its moving speed in 50% and changes the direction to avoid wall. The updated moving speed (*V*_*t*_′) and direction (*D*_*t*_′) at time *t* are defined as follows:

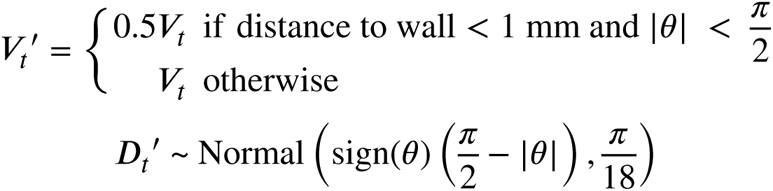

To introduce a temporal correlation in movement speed, we further incorporated a speed inheritance mechanism in the model. The speed at time *t* (*V*_*t*_) is calculated as a weighted sum of 60% of the speed from the previous timestep (*V*_*t*−1_) and 40% of the current adjusted speed (*V*_*t*_′):

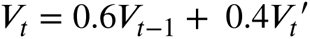

Next, visual stimuli were introduced every 15 s starting 5 min into the experiment, defined by a fixed probability of stopping movement. Immediately after a stimulus is given at time *t*, there’s a 90% chance that the moving speed becomes 0. This state is represented by *S*_*t*_, where *S*_*t*_ = 1 if the moving speed becomes 0, and *S*_*t*_ = 0 otherwise.

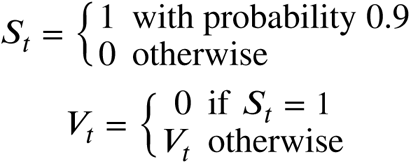

If the moving speed becomes 0 immediately after a stimulus (*S*_*t*_ = 1), the subsequent moving speed for up to 15 s until the next stimulus follows a probability function as below, where *a* and *b* are constants, and *t’* is the time elapsed after the stimulus, up to 15 s.

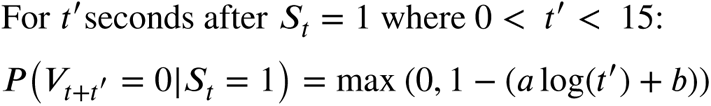

In the single and group conditions, we applied *a* = 0.02 and *b* = 0.92 into the model, reproducing continuous freezing. In the mixed-strain group condition, we simulated 15 strains with different parameters *a* = [0.005, 0.006, 0.007, 0.008, 0.009, 0.01, 0.02, 0.03, 0.04, 0.05, 0.06, 0.07, 0.08, 0.09, 0.1] and *b* = 1.02 − 2.45*a*, based on the experimental data. Lastly, we incorporated the change in motion cues of other individuals from time *t* − 2 to *t* − 1 into individual’s moving speed at time *t*, thereby updating their speed based on social interaction.

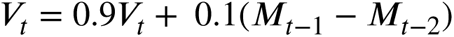

where *M*_*t*−1_−*M*_*t*−2_ represent the change in motion cues between previous time steps. Motion cues were calculated as shown in Fig. 1d, and 10% of the change was incorporated to adjust individual’s speed to others.

## Data availability

Behavioral and population genetic datasets obtained from this study will be available upon publication.

## Code availability

The codes used in this study are available at https://github.com/daikisato12/Sato2024_Fruitfly_collectives.

## Acknowledgements

Computations for behavioral analysis were partially performed on the NIG supercomputer at ROIS National Institute of Genetics. This work was supported by the Japan Society for the Promotion of Science (Grants-in-Aid for Scientific Research JP22K15181 to D.X.S. and JP23H03840 to Y.T.) and by the Sasakawa Scientific Research Grant from The Japan Science Society (to D.X.S.).

## Author contributions

D.X.S. and Y.T. conceived and designed the study. D.X.S. and Y.T. acquired the funding. D.X.S. conducted behavioral experiments with fruit flies, performed agent-based simulation, and analyzed behavioral, single-cell transcriptomic, and population genetic data. D.X.S. and Y.T. wrote and approved the final manuscript.

## Reference

1. Krause, J. & Ruxton, G. D. Living in Groups. (Oxford University Press, 2002).

2. Sumpter, D. J. T. Collective Animal Behavior. (Princeton University Press, 2010).

3. Couzin, I. D. & Krause, J. Self-organization and collective behavior in vertebrates. Adv. Stud. Behav. 32, (2003).

4. Katz, Y., Tunstrøm, K., Ioannou, C. C., Huepe, C. & Couzin, I. D. Inferring the structure and dynamics of interactions in schooling fish. Proceedings of the National Academy of Sciences 108, 18720–18725 (2011).

5. Strandburg-Peshkin, A. et al. Visual sensory networks and effective information transfer in animal groups. Curr. Biol. 23, R709–11 (2013).

6. Bastien, R. & Romanczuk, P. A model of collective behavior based purely on vision. Sci Adv 6, eaay0792 (2020).

7. Davidson, J. D. et al. Collective detection based on visual information in animal groups. J. R. Soc. Interface 18, 20210142 (2021).

8. Williams, H. J. et al. Sensory collectives in natural systems. Elife 12, (2023).

9. Ramdya, P., Schneider, J. & Levine, J. D. The neurogenetics of group behavior in Drosophila melanogaster. J. Exp. Biol. 220, 35–41 (2017).

10. Ferreira, C. H. & Moita, M. A. What can a non-eusocial insect tell us about the neural basis of group behaviour? Curr Opin Insect Sci 36, 118–124 (2019).

11. Couzin-Fuchs, E. & Ayali, A. The social brain of “non-eusocial” insects. Curr. Opin. Insect Sci. 48, 1–7 (2021).

12. Jiang, L. et al. Emergence of social cluster by collective pairwise encounters in Drosophila. eLife 9, e51921 (2020).

13. Zacarias, R., Namiki, S., Card, G. M., Vasconcelos, M. L. & Moita, M. A. Speed dependent descending control of freezing behavior in Drosophila melanogaster. Nat. Commun. 9, 3697 (2018).

14. Ferreira, C. H. & Moita, M. A. Behavioral and neuronal underpinnings of safety in numbers in fruit flies. Nat. Commun. 11, 4182 (2020).

15. Bengochea, M. et al. Numerical discrimination in Drosophila melanogaster. Cell Rep. 42, 112772 (2023).

16. Wice, E. W. & Saltz, J. B. Selection on heritable social network positions is context-dependent in Drosophila melanogaster. Nat. Commun. 12, 1–9 (2021).

17. Tang, W. et al. Genetic control of collective behavior in zebrafish. iScience 23, 100942 (2020).

18. Harpaz, R. et al. Collective behavior emerges from genetically controlled simple behavioral motifs in zebrafish. Sci Adv 7, eabi7460 (2021).

19. Kocher, S. D. et al. The genetic basis of a social polymorphism in halictid bees. Nat. Commun. 9, 4338 (2018).

20. Corral-Lopez, A. et al. Functional convergence of genomic and transcriptomic architecture underlies schooling behaviour in a live-bearing fish. Nature Ecology & Evolution 1–13 (2023).

21. MacKay, T. F. C. et al. The Drosophila melanogaster Genetic Reference Panel. Nature 482, 173–178 (2012).

22. Watanabe, L. P., Gordon, C., Momeni, M. Y. & Riddle, N. C. Genetic networks underlying natural variation in basal and induced activity levels in drosophila melanogaster. G3: Genes, Genomes, Genetics 10, 1247–1260 (2020).

23. Shorter, J. et al. Genetic architecture of natural variation in Drosophila melanogaster aggressive behavior. Proceedings of the National Academy of Sciences 112, E3555– E3563 (2015).

24. Yanagawa, A. et al. Genetic Basis of Natural Variation in Spontaneous Grooming in Drosophila melanogaster. G3: Genes, Genomes, Genetic 10, 3453–3460 (2020).

25. Harbison, S. T. et al. Genome-Wide Association Study of Circadian Behavior in Drosophila melanogaster. Behav. Genet. 49, 60–82 (2019).

26. Jolles, J. W., Boogert, N. J., Sridhar, V. H., Couzin, I. D. & Manica, A. Consistent Individual Differences Drive Collective Behavior and Group Functioning of Schooling Fish. Curr. Biol. 27, 2862–2868.e7 (2017).

27. Jolles, J. W., King, A. J. & Killen, S. S. The Role of Individual Heterogeneity in Collective Animal Behaviour. Trends Ecol. Evol. 35, 278–291 (2020).

28. Takahashi, Y., Tanaka, R., Yamamoto, D., Noriyuki, S. & Kawata, M. Balanced genetic diversity improves population fitness. Proceedings of the Royal Society B: Biological Sciences 285, (2018).

29. Bentzur, A., Ben-shaanan, S., Benishou, J., Costi, E. & Ilany, A. Early Life Experience Shapes Male Behavior and Social Networks in Drosophila Article Early Life Experience Shazpes Male Behavior and Social Networks in Drosophila. Curr. Biol. 1–16 (2021).

30. Nagy, M. et al. Synergistic Benefits of Group Search in Rats. Curr. Biol. 1–6 (2020).

31. Tan, L. et al. Ig Superfamily Ligand and Receptor Pairs Expressed in Synaptic Partners in Drosophila. Cell 163, 1756–1769 (2015).

32. Kurmangaliyev, Y. Z., Yoo, J., Valdes-Aleman, J., Sanfilippo, P. & Zipursky, S. L. Transcriptional Programs of Circuit Assembly in the Drosophila Visual System. Neuron 108, 1045–1057.e6 (2020).

33. Stoks, R., McPeek, M. A. & Mitchell, J. L. Evolution of prey behavior in response to changes in predation regime: damselflies in fish and dragonfly lakes. Evolution 57, 574– 585 (2003).

34. Chelini, M.-C., Willemart, R. H. & Hebets, E. A. Costs and benefits of freezing behaviour in the harvestman Eumesosoma roeweri (Arachnida, Opiliones). Behav. Processes 82, 153–159 (2009).

35. Joesch, M., Schnell, B., Raghu, S. V., Reiff, D. F. & Borst, A. ON and OFF pathways in Drosophila motion vision. Nature 468, 300–304 (2010).

36. Meier, M. et al. Neural circuit components of the Drosophila OFF motion vision pathway. Curr. Biol. 24, 385–392 (2014).

37. Takemura, S.-Y., Lu, Z. & Meinertzhagen, I. A. Synaptic circuits of the Drosophila optic lobe: the input terminals to the medulla. J. Comp. Neurol. 509, 493–513 (2008).

38. Borst, A. & Groschner, L. N. How Flies See Motion. Annu. Rev. Neurosci. 46, 17–37 (2023).

39. Borst, A. & Helmstaedter, M. Common circuit design in fly and mammalian motion vision. Nat. Neurosci. 18, 1067–1076 (2015).

40. Sanes, J. R. & Zipursky, S. L. Synaptic Specificity, Recognition Molecules, and Assembly of Neural Circuits. Cell 181, 536–556 (2020).

41. Simon, F. & Konstantinides, N. Single-cell transcriptomics in the Drosophila visual system: Advances and perspectives on cell identity regulation, connectivity, and neuronal diversity evolution. Dev. Biol. 479, 107–122 (2021).

42. Özel, M. N. et al. Neuronal diversity and convergence in a visual system developmental atlas. Nature 589, 88–95 (2021).

43. Pak, W. L., Grossfield, J. & Arnold, K. S. Mutants of the visual pathway of Drosophila melanogaster. Nature 227, 518–520 (1970).

44. Sridhar, V. H., Roche, D. G. & Gingins, S. Tracktor: Image-based automated tracking of animal movement and behaviour. Methods Ecol. Evol. 10, 815–820 (2019).

45. Singh, M., Ceccarelli, S. & Hamblin, J. Estimation of heritability from varietal trials data. Theor. Appl. Genet. 86, 437–441 (1993).

46. Kruijer, W. et al. Marker-based estimation of heritability in immortal populations. Genetics 199, 379–398 (2015).

47. Chang, C. C. et al. Second-generation PLINK: rising to the challenge of larger and richer datasets. Gigascience 4, 7 (2015).

48. Wu, Y. et al. Fast estimation of genetic correlation for biobank-scale data. Am. J. Hum. Genet. 109, 24–32 (2022).

49. Jiang, L. et al. A resource-efficient tool for mixed model association analysis of large-scale data. Nat. Genet. 51, 1749–1755 (2019).

50. Danecek, P. et al. The variant call format and VCFtools. Bioinformatics 27, 2156–2158 (2011).

