## Supplementary Tables and Figures for "Neurogenomic diversity enhances collective antipredator performance in *Drosophila*"

**Supplementary information**

This file includes:

- Tables S1
- Figure S1–10

Table S1. Fly strains used in this study.

| BDSC # | Genotype | Name |
|---|---|---|
| 9047 | w[*] norpA[P12] | *norpA* |
| 28838 | w[*]; P{w[+mC]=UAS-TeTxLC.tnt}G2 | *UAS-TNT* |
| 28840 | w[*]; P{w[+mC]=UAS-TeTxLC.(-)V}A2 | *UAS-IMPTNT* |
| 25840 | y[1] v[1]; P{y[+t7.7] v[+t1.8]=TRiP.JF01858}attP2 | *UAS-Ptp99A*$^{RNAi}$ |
| 41369 | w[1118]; P{y[+t7.7] w[+mC]=GMR9B08-GAL4}attP2/TM3, Sb[1] | *R9B08-GAL4* |
| 24843 | w[1118]; Mi{GFP[E.3xP3]=ET1}Ptp99A[MB04947] | *Ptp99A-GAL4* |
| 28122 | NA | DGRP-21 |
| 28123 | NA | DGRP-26 |
| 28124 | NA | DGRP-28 |
| 28126 | NA | DGRP-41 |
| 28128 | NA | DGRP-45 |
| 29652 | NA | DGRP-57 |
| 28130 | NA | DGRP-69 |
| 28131 | NA | DGRP-73 |
| 28132 | NA | DGRP-75 |
| 28135 | NA | DGRP-88 |
| 28137 | NA | DGRP-93 |
| 28138 | NA | DGRP-101 |
| 28139 | NA | DGRP-105 |
| 28140 | NA | DGRP-109 |
| 28141 | NA | DGRP-129 |
| 28142 | NA | DGRP-136 |
| 28144 | NA | DGRP-142 |
| 28145 | NA | DGRP-149 |
| 28146 | NA | DGRP-153 |
| 28148 | NA | DGRP-161 |
| 28149 | NA | DGRP-176 |
| 28151 | NA | DGRP-181 |
| 28152 | NA | DGRP-189 |
| 28153 | NA | DGRP-195 |
| 25174 | NA | DGRP-208 |
| 28154 | NA | DGRP-217 |
| 28157 | NA | DGRP-228 |
| 28275 | NA | DGRP-235 |
| 28164 | NA | DGRP-280 |
| 28165 | NA | DGRP-287 |

| | | |
|---|---|---|
| 25175 | NA | DGRP-301 |
| 25176 | NA | DGRP-303 |
| 28166 | NA | DGRP-309 |
| 25181 | NA | DGRP-315 |
| 28168 | NA | DGRP-318 |
| 29654 | NA | DGRP-320 |
| 25182 | NA | DGRP-324 |
| 25183 | NA | DGRP-335 |
| 28174 | NA | DGRP-340 |
| 25184 | NA | DGRP-357 |
| 25185 | NA | DGRP-358 |
| 25186 | NA | DGRP-360 |
| 25187 | NA | DGRP-362 |
| 25445 | NA | DGRP-365 |
| 28182 | NA | DGRP-370 |
| 28184 | NA | DGRP-373 |
| 25188 | NA | DGRP-375 |
| 25189 | NA | DGRP-379 |
| 28189 | NA | DGRP-382 |
| 28191 | NA | DGRP-385 |
| 25192 | NA | DGRP-399 |
| 29656 | NA | DGRP-405 |
| 25193 | NA | DGRP-427 |
| 25194 | NA | DGRP-437 |
| 28199 | NA | DGRP-443 |
| 28200 | NA | DGRP-461 |
| 25195 | NA | DGRP-486 |
| 28204 | NA | DGRP-502 |
| 28205 | NA | DGRP-508 |
| 29659 | NA | DGRP-513 |
| 25197 | NA | DGRP-517 |
| 28207 | NA | DGRP-531 |
| 28208 | NA | DGRP-535 |
| 25198 | NA | DGRP-555 |
| 28211 | NA | DGRP-563 |
| 28212 | NA | DGRP-584 |
| 28213 | NA | DGRP-589 |
| 28218 | NA | DGRP-703 |
| 25200 | NA | DGRP-707 |
| 25201 | NA | DGRP-712 |

| | | |
|---|---|---|
| 28219 | NA | DGRP-716 |
| 28220 | NA | DGRP-721 |
| 25202 | NA | DGRP-730 |
| 25203 | NA | DGRP-732 |
| 28224 | NA | DGRP-748 |
| 28226 | NA | DGRP-757 |
| 28227 | NA | DGRP-761 |
| 25204 | NA | DGRP-765 |
| 25205 | NA | DGRP-774 |
| 25206 | NA | DGRP-786 |
| 28233 | NA | DGRP-796 |
| 25207 | NA | DGRP-799 |
| 28234 | NA | DGRP-801 |
| 28236 | NA | DGRP-804 |
| 28238 | NA | DGRP-808 |
| 28239 | NA | DGRP-810 |
| 25208 | NA | DGRP-820 |
| 28244 | NA | DGRP-822 |
| 28245 | NA | DGRP-832 |
| 28246 | NA | DGRP-837 |
| 28247 | NA | DGRP-843 |
| 28248 | NA | DGRP-849 |
| 25209 | NA | DGRP-852 |
| 28251 | NA | DGRP-855 |
| 28252 | NA | DGRP-857 |
| 25210 | NA | DGRP-859 |
| 28254 | NA | DGRP-879 |
| 28256 | NA | DGRP-884 |
| 28257 | NA | DGRP-890 |
| 28258 | NA | DGRP-892 |
| 28260 | NA | DGRP-897 |
| 28262 | NA | DGRP-907 |
| 28264 | NA | DGRP-911 |
| 28265 | NA | DGRP-913 |

7

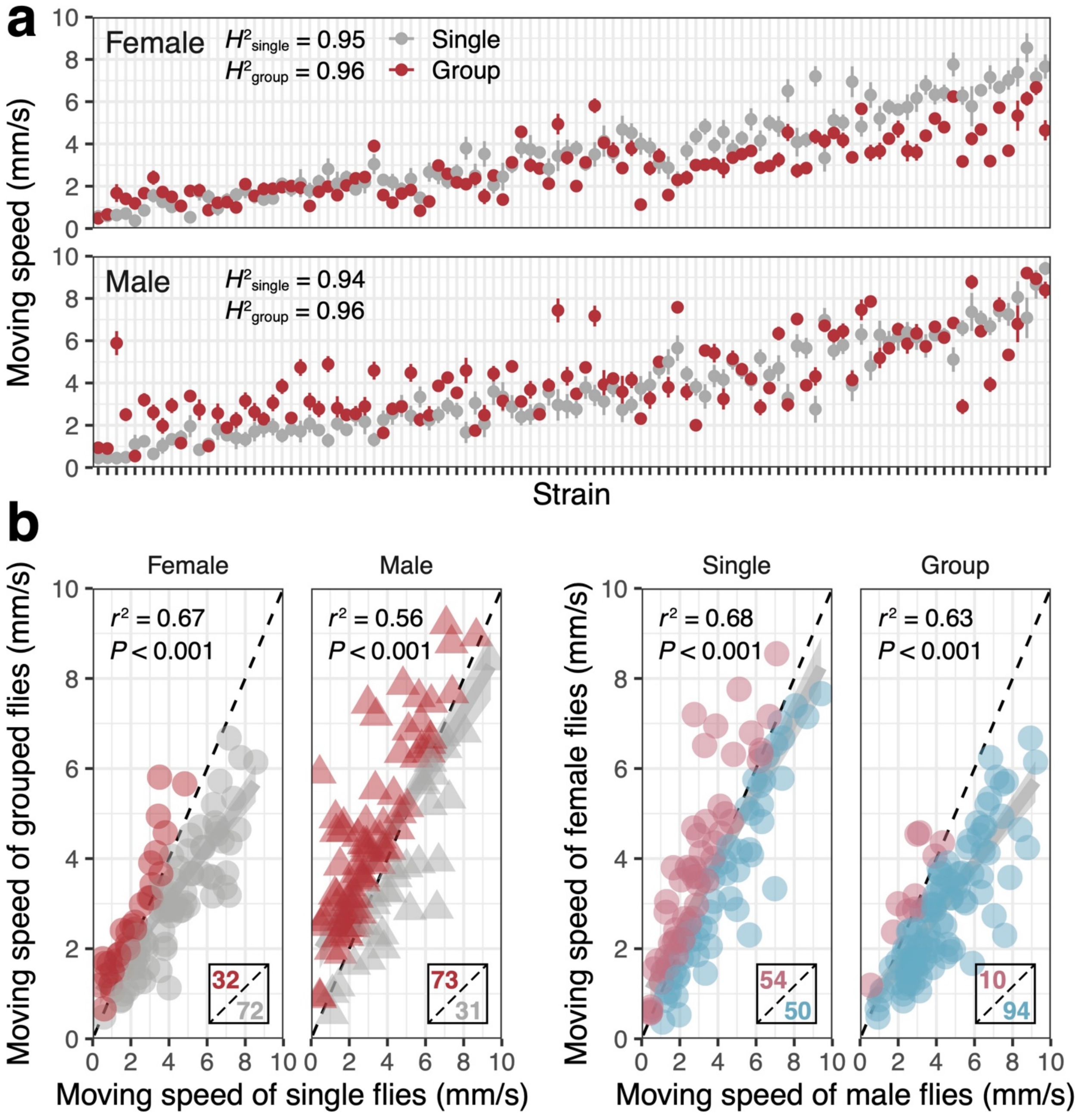


**Figure S1. Inter-strain variation in average moving speed. (a)** Mean moving speed of DGRP strains. Color corresponds to social conditions (red for group and grey for single conditions), and the error bar represents standard errors. **(b)** The correlations in the mean moving speed between social conditions and sexes. The points beneath or above the dashed lines, indicating y = x, are highlighted in different colors, and their counts are described in subpanels below.

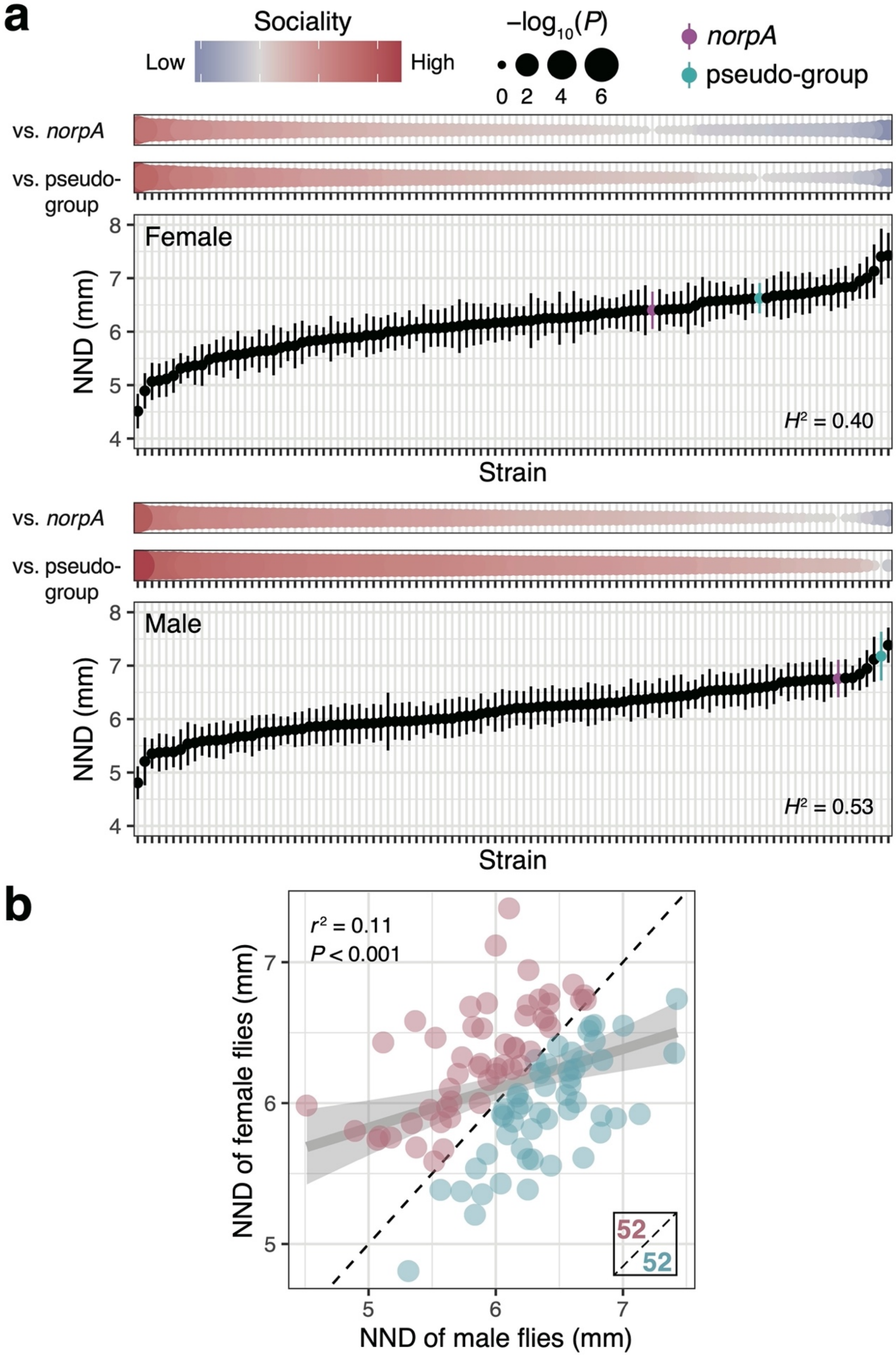

a
Sociality
Low
High
−log10(P)
0 2 4 6
norpA
pseudo-group
vs. norpA
vs. pseudo-group
Female
NND (mm)
H² = 0.40
Strain
vs. norpA
vs. pseudo-group
Male
NND (mm)
H² = 0.53
Strain
b
r² = 0.11
P < 0.001
NND of female flies (mm)
NND of male flies (mm)
52
52

**Figure S2. Inter-strain variation in nearest neighbor distance (NND).** Mean nearest neighbor distance (NND) for DGRP strains. **(a)** *NorpA* mutants and pseudo-group (virtual combinations of single flies) data are colored in magenta and light blue, respectively, and were used as controls with no social preference. The difference in NND with the control groups was evaluated by the least squares means, and the $\log_{10}$-transformed uncorrected *P*-values, indicated in the size of points, are shown on the top panels with estimates of the difference shown in the heatmap, as a measure of sociality. The error bar represents standard errors. **(b)** The correlations in the mean NND between sexes. The points beneath or above the dashed lines, indicating y = x, are highlighted in different colors, and their counts are described in subpanels below.

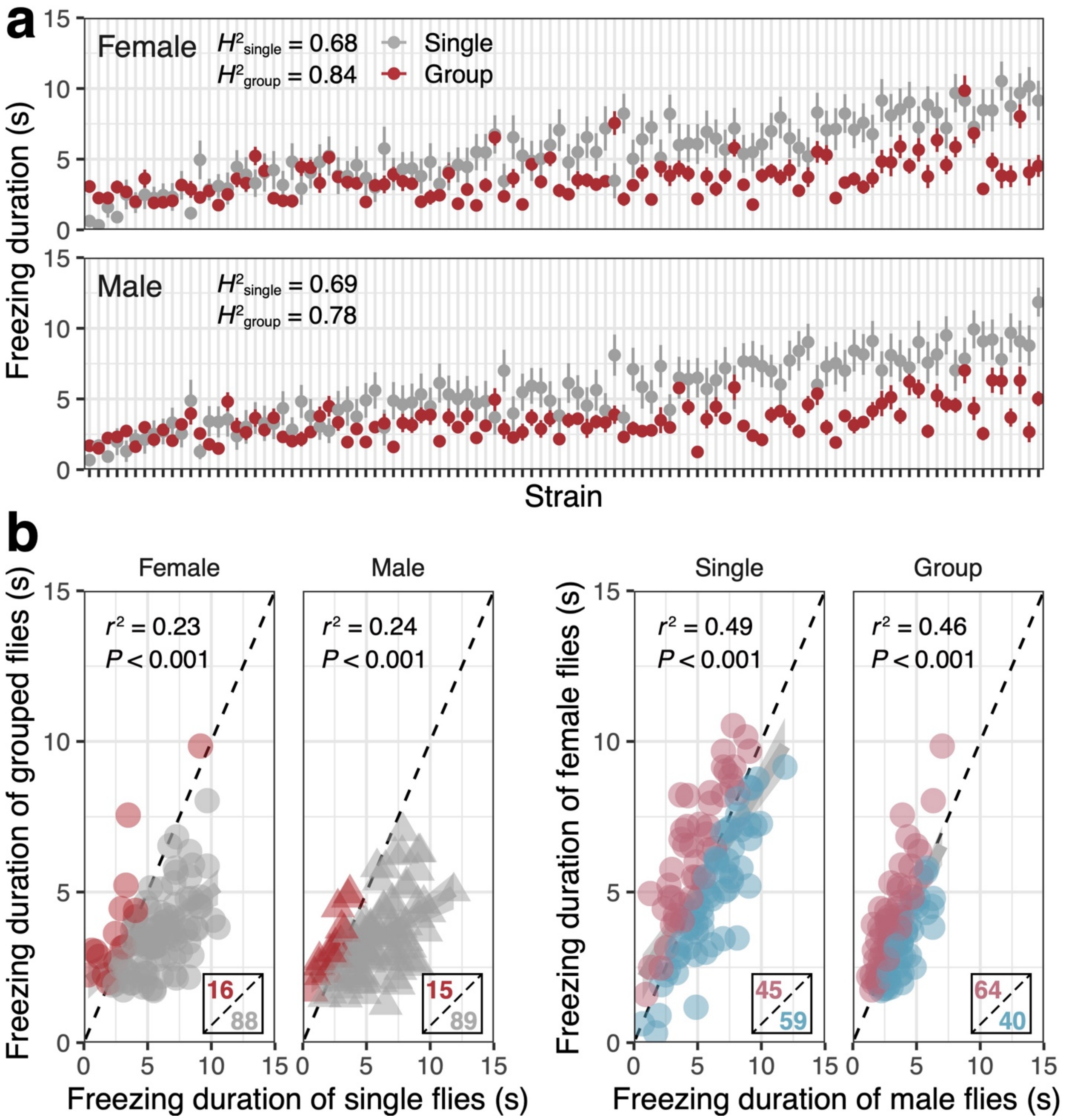


**Figure S3. Inter-strain variation in freezing duration. (a)** Mean freezing duration of DGRP strains. Color corresponds to social conditions (red for group and grey for single conditions), and the error bar represents standard errors. **(b)** The correlations in the mean freezing duration between social conditions and sexes. The points beneath or above the dashed lines, indicating y = x, are highlighted in different colors, and their counts are described in subpanels below.

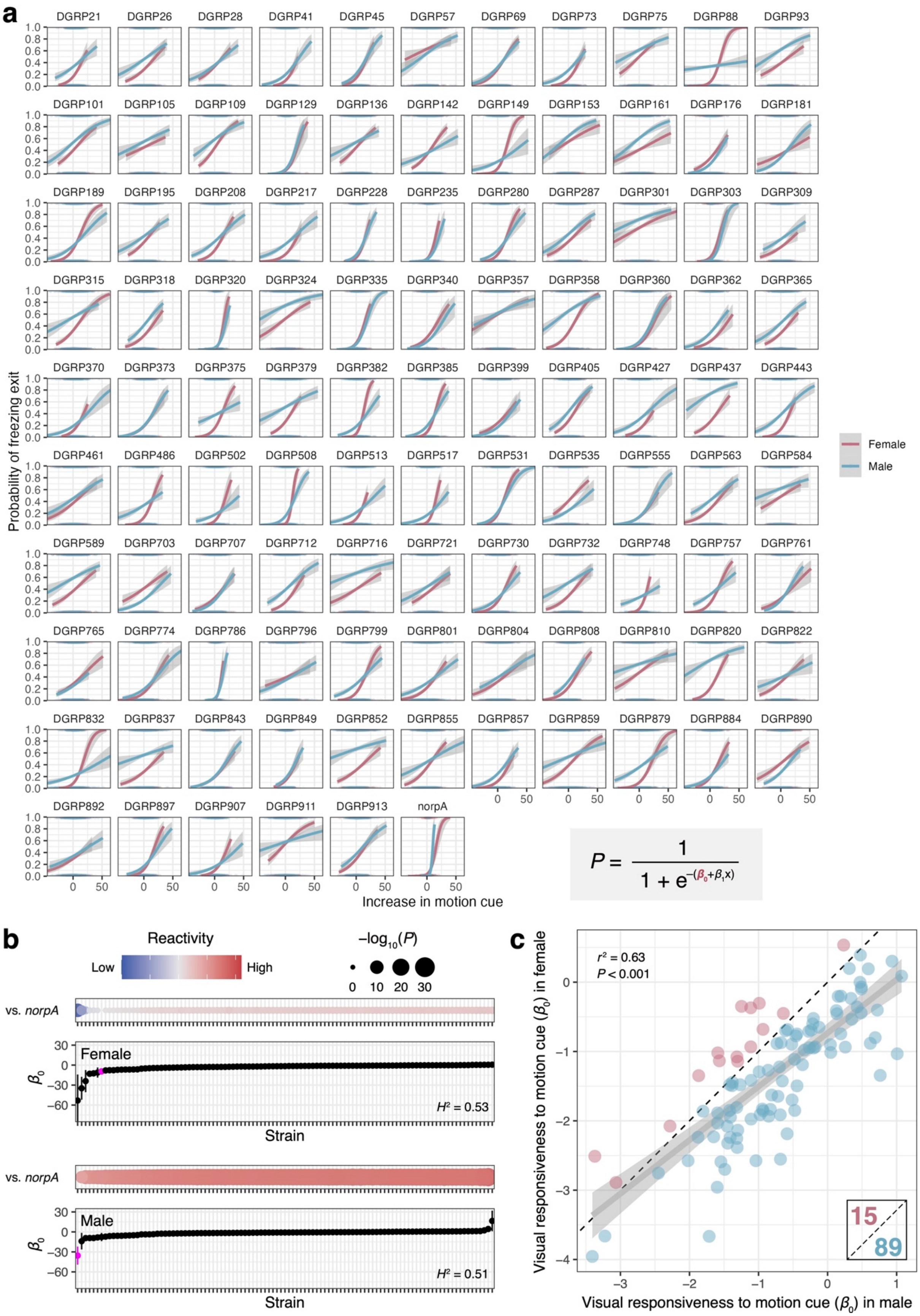
a
DGRP21 DGRP26 DGRP28 DGRP41 DGRP45 DGRP57 DGRP69 DGRP73 DGRP75 DGRP88 DGRP93
DGRP101 DGRP105 DGRP109 DGRP129 DGRP136 DGRP142 DGRP149 DGRP153 DGRP161 DGRP176 DGRP181
DGRP189 DGRP195 DGRP208 DGRP217 DGRP228 DGRP235 DGRP280 DGRP287 DGRP301 DGRP303 DGRP309
DGRP315 DGRP318 DGRP320 DGRP324 DGRP335 DGRP340 DGRP357 DGRP358 DGRP360 DGRP362 DGRP365
DGRP370 DGRP373 DGRP375 DGRP379 DGRP382 DGRP385 DGRP399 DGRP405 DGRP427 DGRP437 DGRP443
DGRP461 DGRP486 DGRP502 DGRP508 DGRP513 DGRP517 DGRP531 DGRP535 DGRP555 DGRP563 DGRP584
DGRP589 DGRP703 DGRP707 DGRP712 DGRP716 DGRP721 DGRP730 DGRP732 DGRP748 DGRP757 DGRP761
DGRP765 DGRP774 DGRP786 DGRP796 DGRP799 DGRP801 DGRP804 DGRP808 DGRP810 DGRP820 DGRP822
DGRP832 DGRP837 DGRP843 DGRP849 DGRP852 DGRP855 DGRP857 DGRP859 DGRP879 DGRP884 DGRP890
DGRP892 DGRP897 DGRP907 DGRP911 DGRP913 norpA
Probability of freezing exit
Increase in motion cue
Female
Male
$P = \frac{1}{1 + e^{-(\beta_0 + \beta_1 x)}}$
b
Reactivity
Low
High
$-\log_{10}(P)$
0 10 20 30
vs. norpA
Female
$\beta_0$
$H^2 = 0.53$
Strain
vs. norpA
Male
$\beta_0$
$H^2 = 0.51$
Strain
c
$r^2 = 0.63$
$P < 0.001$
15
89
Visual responsiveness to motion cue ($\beta_0$) in female
Visual responsiveness to motion cue ($\beta_0$) in male

**Figure S4. Inter-strain variation in the responsiveness to other individuals' movement. (a)** The logistic curve representing reaction toward motion cues of other conspecifics of DGRP strains. Given the logistic formula shown on the right bottom, we estimated and used the intercept $\beta_0$ as an index of visual responsiveness of flies toward conspecifics' motion cues. **(b)** Mean visual responsiveness ($\beta_0$) for DGRP strains. *NorpA* mutants are colored in magenta and were used as controls with visual defects. The difference in $\beta_0$with the control groups was evaluated by the least squares means, and the $\log_{10}$-transformed uncorrected *P*-values, indicated in the size of points, are shown on the top panels with estimates of the difference shown in the heatmap, as a measure of visual responsiveness. The error bar represents standard errors. **(c)** The correlations in the $\beta_0$ between sexes. The points beneath or above the dashed lines, indicating y = x, are highlighted in different colors, and their counts are described in subpanels below.

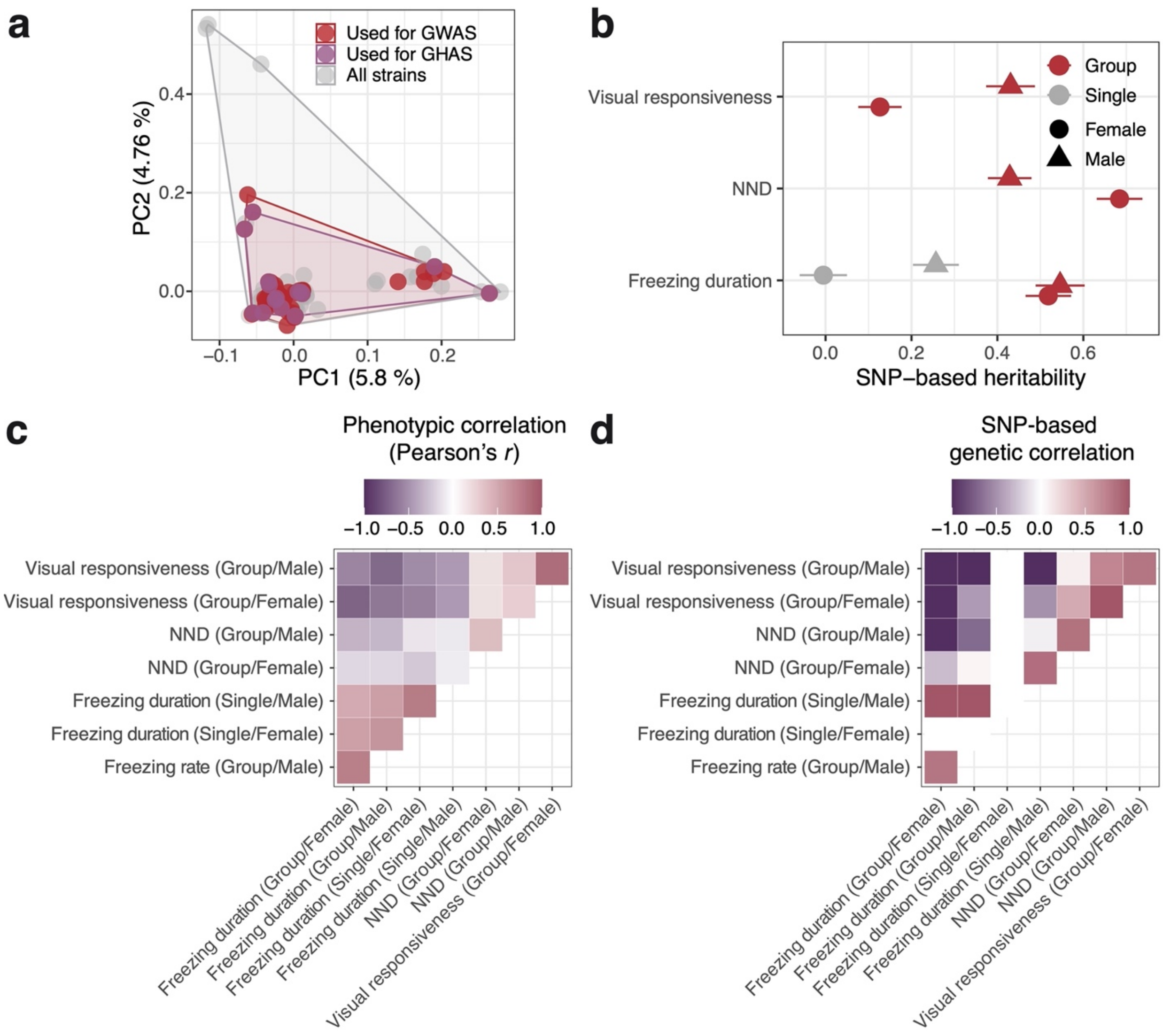


**Figure S5. Genetic distances among DGRP strains and heritability estimates and genetic correlation among analyzed phenotypes used in our study. (a)** Principal component analysis for a whole set of genotypes of 4,438,427 SNPs from 205 strains of DGRP showed that 15 strains used in mixed-group experiments (colored in purple) were as genetically diverse as the 104 strains used in GWAS (colored in red). **(b)** SNP-based heritability of the analyzed phenotypes estimated from genomic and phenotypic data. The error bar represents standard errors. The heritability of female freezing duration in the single condition was estimated to be negative, indicating that the calculation could not be converged possibly due to the lack of sufficient additive genetic effects. **(c)** Phenotypic and **(d)** SNP-based genetic correlation among the phenotypes, except for the female freezing duration in the single condition.

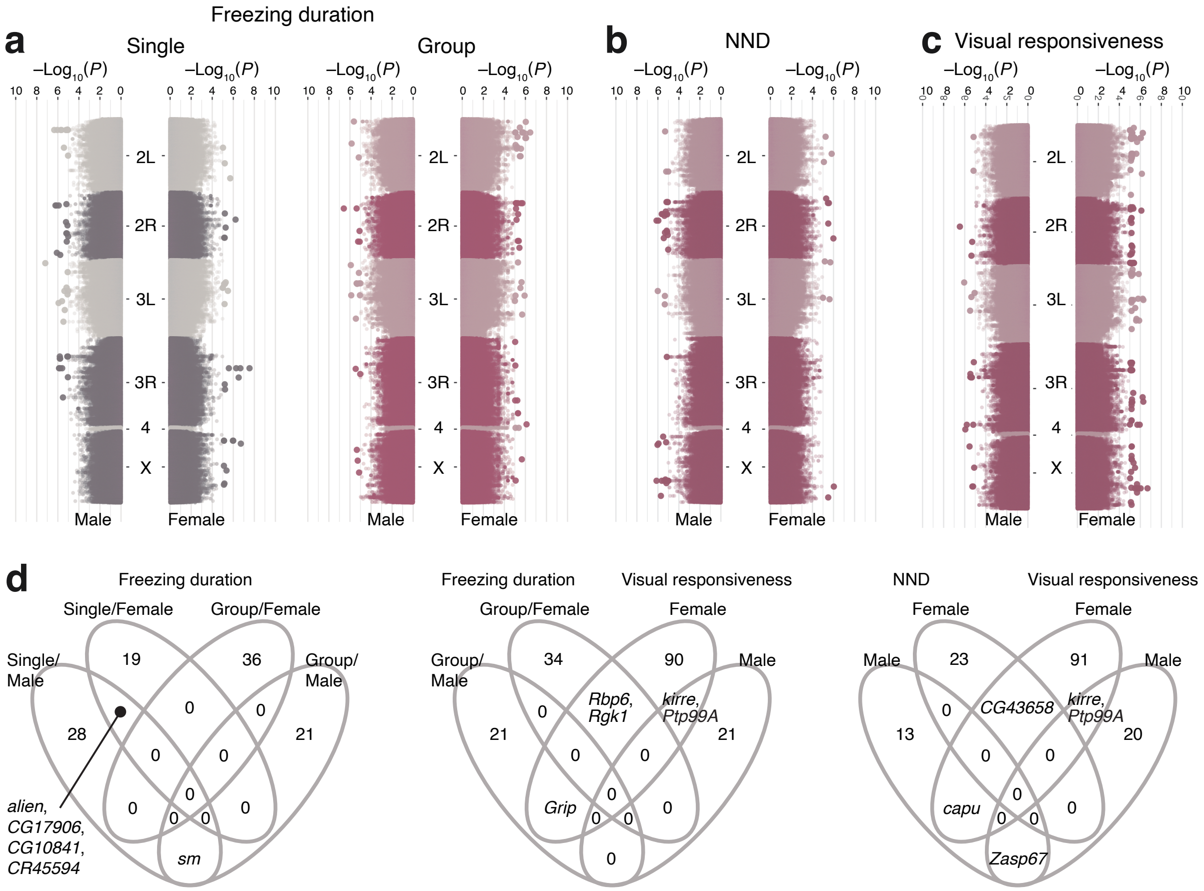


**Figure S6. Manhattan plots and genes detected in GWAA for freezing duration, NND, and visual responsiveness.** Manhattan plots for **(a)** freezing duration, **(b)** NND, and **(c)** visual responsiveness. Colors represent different chromosomes within a panel, and the size of points indicates statistical significance ($P < 1.0 \times 10^{-5}$). **(d)** Overlap of genes adjacently located to detected loci among traits and sexes are shown in Venn diagrams.

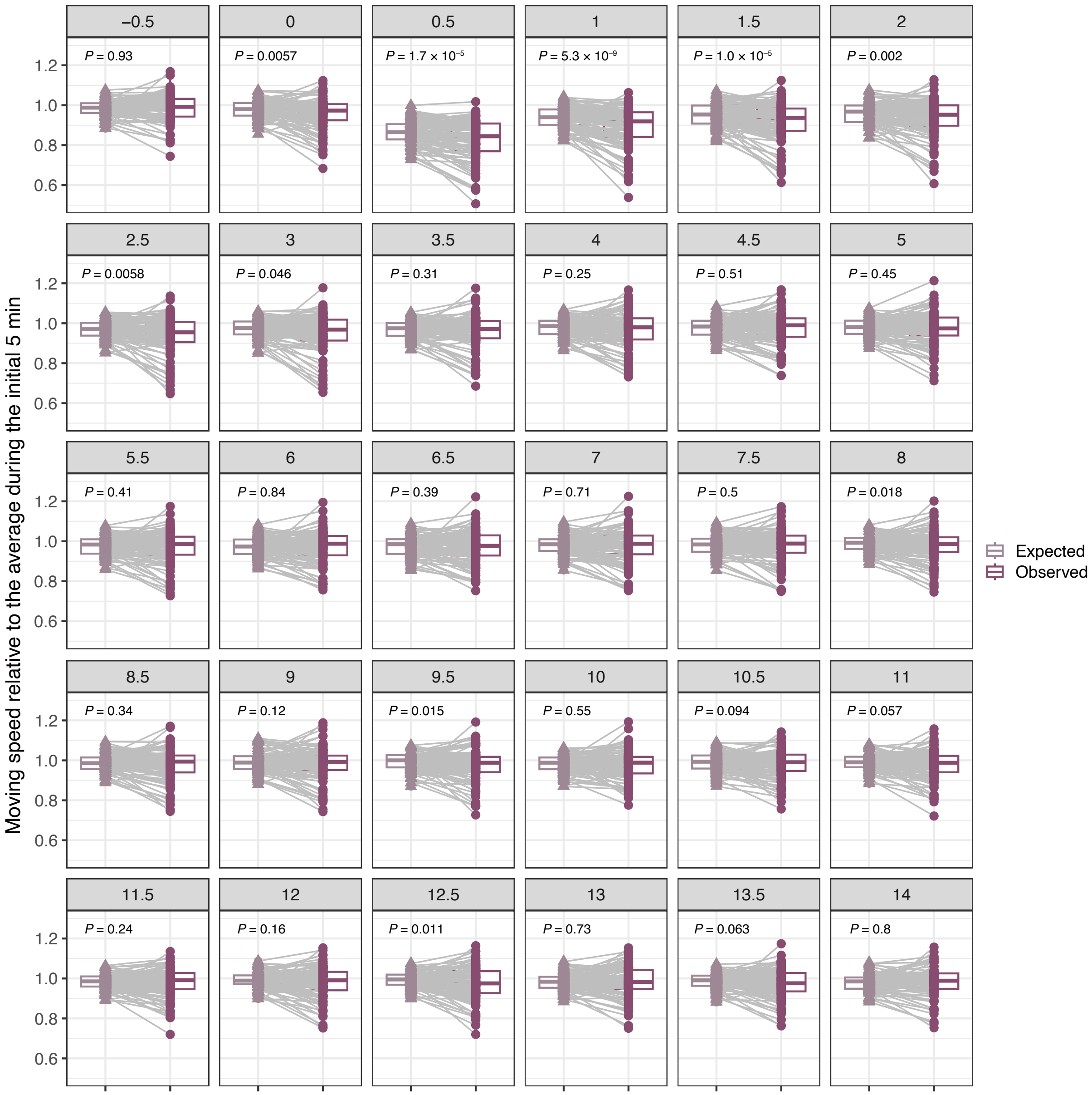


**Figure S7. Mean relative moving speed decreased in mixed-strain groups at immediately after looming stimulus.** Expected and observed values of mean relative moving speed of mixed-strain groups is shown for each 0.5 s bin after stimulus. Statistical significance was evaluated by Wilcoxon's signed rank test.

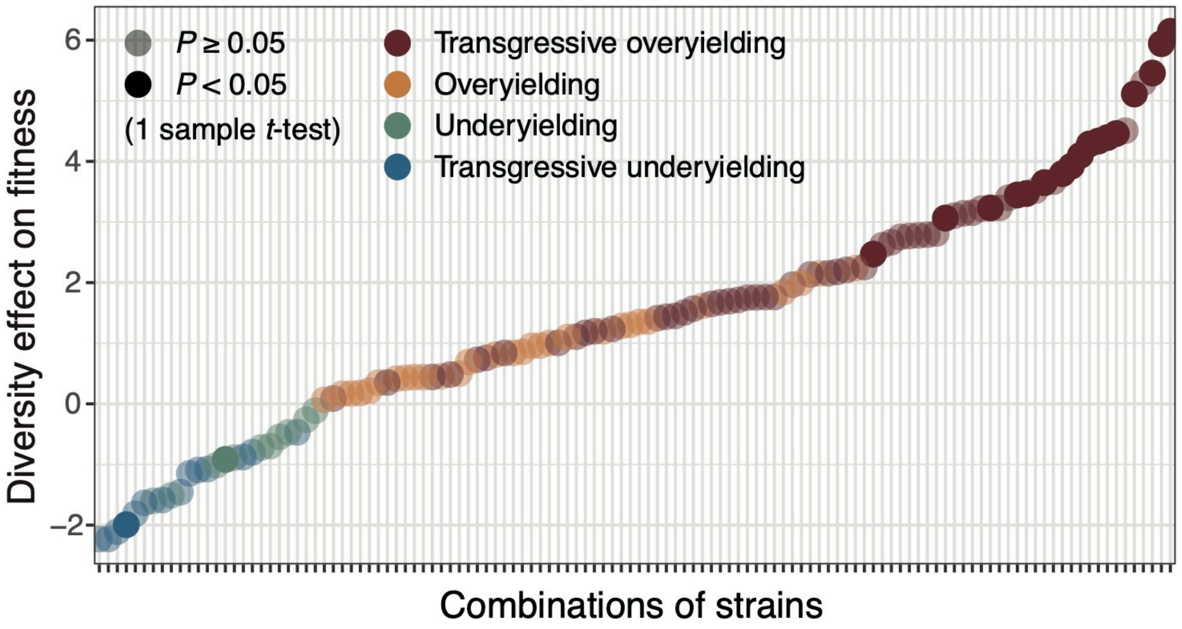


**Figure S8. Diversity effect on virtual fitness measured in total 105 combinations of pairs of strains.** Diversity effect, defined as the difference between expected and observed values of virtual fitness (see Materials and Methods) are sorted along combinations of used strains in mixed-group experiments. Overyielding and underyielding (see Fig. 4d) are colored in different color, and statistical significance is indicated in transparency of plots. One sample *t*-test was used to test statistical significance for the deviation of diversity effect from 0.

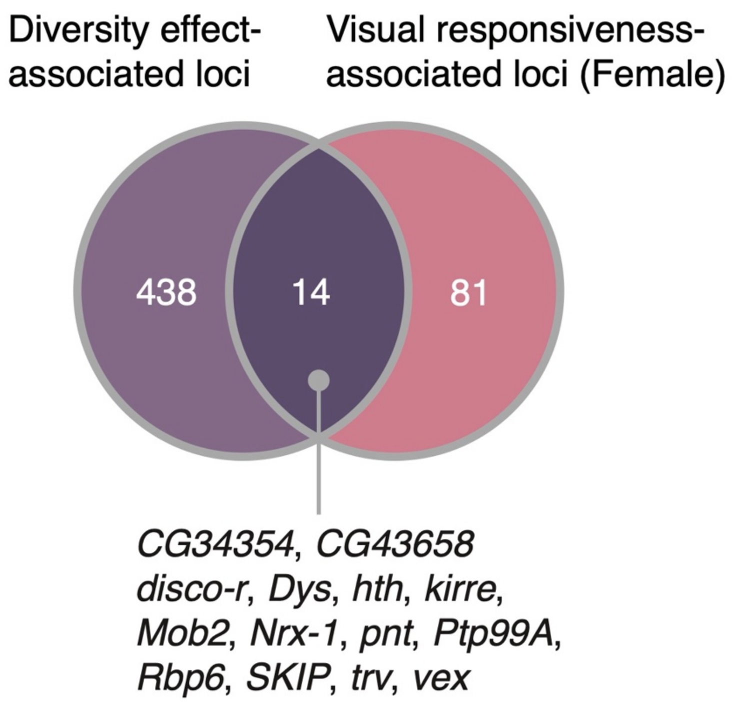


**Figure S9. Venn diagram showing the overlap of loci detected in GWAS and GHAS.** Overlapped genes detected in both GWAS for visual responsiveness in females and GHAS includes *kirre* and *Ptp99A*, indicating the contribution of genetic diversity involved in visual neuron development to emergent group dynamics.

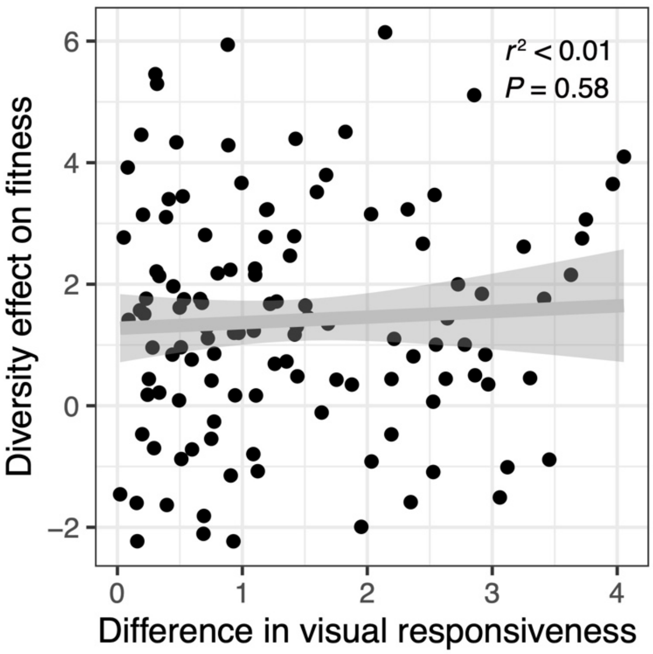


**Figure S10. Correlation between inter-strain difference in visual responsiveness and diversity effect in virtual fitness observed in mixed-strain group experiments.** Unlike the freezing duration (see Fig. 4d), difference in visual responsiveness between strains is not correlated with achieved diversity effect.
